# Comparative genomics for the elucidation of multidrug resistance (MDR) in *Candida lusitaniae*

**DOI:** 10.1101/780833

**Authors:** Abhilash Kannan, Sandra A. Asner, Emilie Trachsel, Steve Kelly, Josie Parker, Dominique Sanglard

## Abstract

Multidrug resistance (MDR) has emerged in hospitals due to the use of several agents administered in combination or sequentially to the same individual. We reported earlier MDR in *Candida lusitaniae* during therapy with amphotericin B (AmB), azoles and candins. We used here comparative genomic approaches between the initial susceptible isolate and 4 other isolates with different MDR profiles. From a total of 18 non-synonymous SNPs (NSS) in genome comparisons with the initial isolate, six could be associated with MDR. One of the SNPs occurred in a putative transcriptional activator (*MRR1*) resulting in a V668G substitution in isolates resistant to azoles and 5-fluorocytosine (5-FC). We demonstrated by gene editing that *MMR1* acted by upregulation of *MFS7* (a multidrug transporter) in the presence of the V668G substitution. *MFS7* itself mediated not only azole resistance but also 5-FC resistance, which represents a novel resistance mechanism for this drug class. Three other distinct NSS occurred in *FKS1* (a glucan synthase that is targeted by candins) in three candin-resistant isolates. Lastly, two other NSS in *ERG3* and *ERG4* (ergosterol biosynthesis) resulting in non-sense mutations were revealed in AmB-resistant isolates, one of which accumulated the 2 *ERG* NSS. AmB-resistant isolates lacked ergosterol and exhibited sterol profiles consistent with *ERG3* and *ERG4* defects. In conclusion, this genome analysis combined with genetics and metabolomics helped to decipher the resistance profiles identified in this clinical case. MDR isolates accumulated 6 different mutations conferring resistance to all antifungal agents used in medicine. This case study illustrates the capacity of *C. lusitaniae* to rapidly adapt under drug pressure within the host.

**Importance:** Antifungal resistance is an inevitable phenomenon when fungal pathogens get exposed to antifungal drugs. These drugs can be grouped in 4 distinct classes (azoles, candins, polyenes, pyrimidine analogs) and are used in different clinical settings. Failures in therapy implicates the sequential or combined use of these different drug classes, which can result in some cases in the development of multidrug resistance (MDR). MDR is particularly challenging in the clinic since it drastically reduces possible treatment alternatives. In this study, we report the rapid development of MDR in *Candida lusitaniae* in a patient, which became resistant to all known antifungal agents used up to now in medicine. To understand how MDR developed in *C. lusitaniae*, whole genome sequencing followed by comparative genome analysis was undertaken in sequential MDR isolates. This helped to detect all specific mutations linked to drug resistance and explained the different MDR patterns exhibited by the clinical isolates.

## Introduction

*Candida* (teleomorph *Clavispora*) *lusitaniae* is a ubiquitous ascomyceteous yeast that can be recovered from soils and water, but can survive in different hosts (birds, mammals). It is known an opportunistic haploid yeast and is the cause of infrequent candidemia (1). Mortality due to *C. lusitaniae* fungemia varies extensively (5 to 50%) and has often been associated with resistance to amphotericin B (Amb) (2). Even if *C. lusitaniae* is considered as susceptible to most systemic antifungal agents, several reports have documented development of antifungal resistance. Usually, antifungal resistance is restricted to a single agent. For example, Desnos *et al.* (3) reported candin resistance in *C. lusitaniae* from a patient treated with caspofungin over 11-17 days. Resistance was associated with a missense mutation (S645F) in the *FKS1* gene encoding β-1,3 glucan synthase, the target of candins in several fungal species. Flucytosine resistance has also been described in *C. lusitaniae* and is associated with defects in cytosine permease (4). Recently, *C. lusitaniae* resistance to fluconazole was reported in the lungs of cystic fibrosis patients, even in the absence of a specific treatment with this drug (5). Simultaneous resistance to AmB and fluconazole was described in a patient treated intermittently with the two agents (6). We reported that *C. lusitaniae* can develop multidrug resistance (MDR), which is understood as resistance to at least 2 different drug classes. MDR occurred in 4 distinct *C. lusitaniae* isolates recovered from a patient treated sequentially with different antifungal drugs including azoles, AmB and candins (7). Antifungal pressure selected distinct MDR profiles that were corresponding to the administered antifungals. The systemic sequential isolates that were obtained from the treated patient were showed to be related with each other suggesting that they had a common resistance patterns (7) and only three separate mutations in *FKS1* resulting in candin resistance were identified. Here we applied whole genome sequencing approaches in the sequential isolates of the treated patient and compared them with the most susceptible isolate that was recovered at early stages of therapy. Genome comparisons revealed several genome alterations explaining the different antifungal resistance profiles of the clinical strains and genetic approaches confirmed their roles in antifungal resistance.

## RESULTS

### C. lusitaniae genomes

We determined earlier that the *C. lusitaniae* sequential isolates (named P1 to P5) recovered from the treated patient were related with each other using restriction sites polymorphisms approaches. Isolate P1 was the earliest but still drug-susceptible isolates, while the other (P4-P5) exhibited MDR (7). To enable complete comparisons between these strains, we carried out their genome sequencing using the Pacbio technology to produce large telomere to telomere assemblies. Five independent assemblies were obtained with the 5 isolates containing each 8 major contigs (smaller contigs corresponding to mitochondrial genomes were ignored). These 8 major contigs were likely to correspond to the 8 suggested chromosomes of *C. lusitaniae* previously described either by supercontigs assemblies of *C. lusitaniae* ATCC 42720 (8), or by chromosome separation from electrophoresis (9) and centromere mapping (10). These Pacbio contigs were designated as Chr 1 to Chr 8 and were sorted by their sizes (Table S3). We first compared the genomes of strain P1 with ATCC 42720 by whole genome alignments. Several chromosome re-arrangements were observed between the two genomes (Fig. 1A), but principally occurring at chromosome ends between Chr 2 and Chr 6 and between Chr 3 and Chr 4. Genome alignments between isolates P1 to P5 did not reveal major re-arrangements, except for a terminal chromosome fragment of Chr 6, which was positioned at the opposite chromosome ends in isolates P2 and P4 (Fig. 1B). In addition, P2 and P5 isolates exhibited a sequence gap at the left side of Chr 4 (Fig. 1B), which was the result of a 30 kb fragment duplication in isolates P1, P3 and P4 (see below).

**Figure 1:**
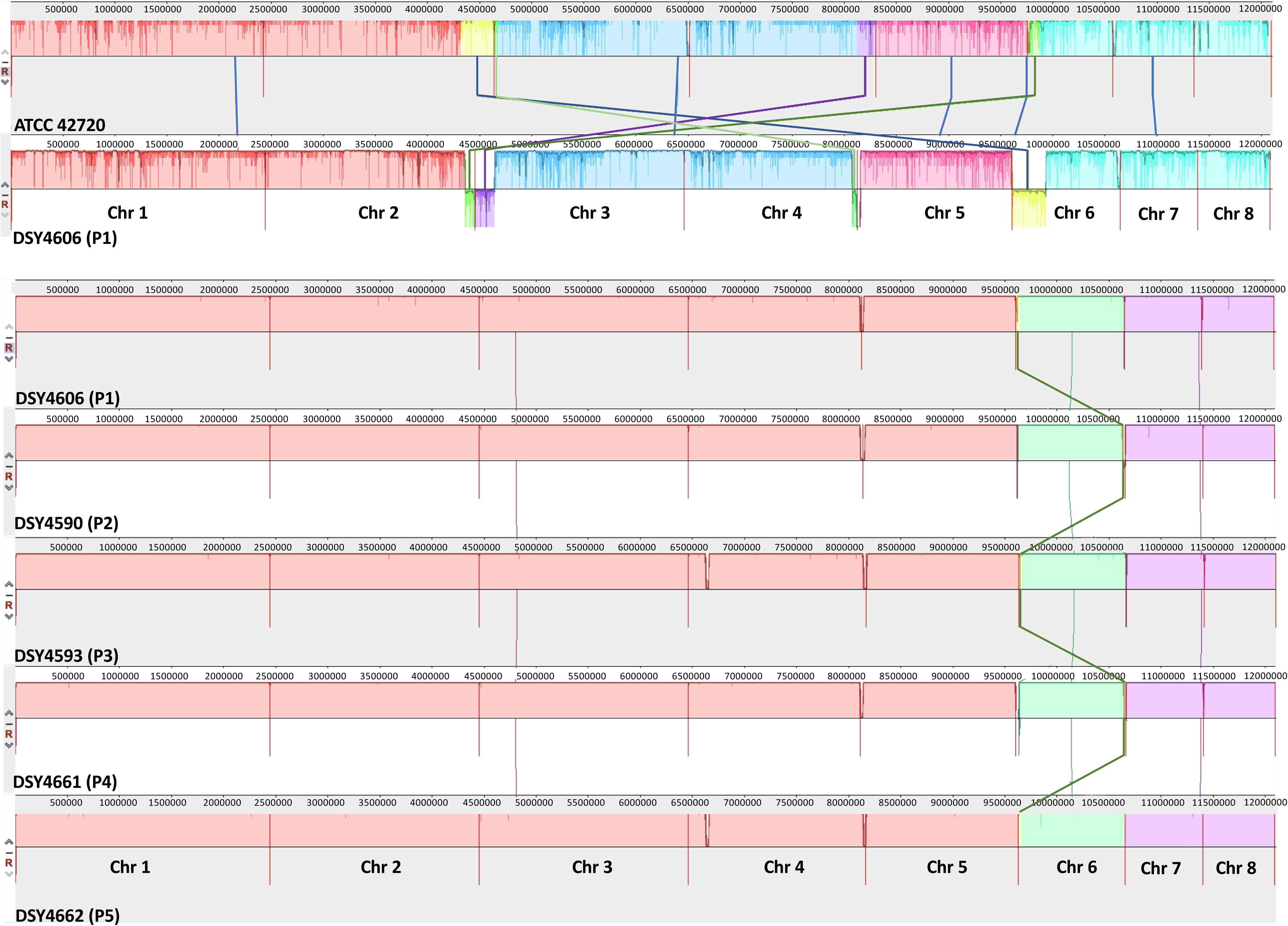
Whole genome alignments. Panel A: alignment between ATCC 42720 and isolate P1. Panel B: alignment between isolates P1 to P5. Alignments were obtained using Mauve (version 2015-02-25) with default parameters and aligner software Muscle 3.6. Vertical red bars indicate separations between the 8 chromosomes. Colored cross-lines between chromosomes of individual isolates indicate major translocations. Isolate designations are both given as laboratory collection and as earlier published in parentheses (7). Color intensities between chromosomes indicate SNP densities between comparisons.

To allow further genome comparisons, genome data of isolate P1 was subsequently subjected to gene annotations (see Material and Methods). This annotation served as a basis for P2 to P5 genome annotations. Genome characteristics of P1 to P5 are supplied in Table 1. The number of detected CDS were ranging from 5676 to 5684 in P1 to P5 genomes. These numbers lie within ranges estimated in genomes of ATCC 42720 and CBS 6936 (Table 1). The genome of CBS 6936 was recently reported but did not reach the level of whole chromosome assemblies (11). Given that P1 and ATCC 42720 genomes were chromosome assemblies, these two genomes were systematically compared for the occurrence of SNPs resulting in synonymous and non-synonymous changes in the encoded proteins (supplementary File S1). Our data suggested the existence of 83225 SNPs between the CDS of these two genomes. The count of non-CDS SNPs was up to 118174 between the two strains, which results in a SNP density of 1 about SNP per 60 bp. This density is higher than observed between *Candida* isolates of the same species (8). A density plot of the detected SNPs along the chromosomes revealed regions of high SNP densities separated by stretches of low densities which are indicative of recombination (Fig. S1). *C. lusitaniae* is one of the few *Candida* spp that is able to undergo mating and meiosis and therefore it can be anticipated that recombination events could be traced to some extent (12). Similar conclusions deduced from SNP density plots were drawn by genome comparisons of *C. glabrata* genomes (13). P1 is of alpha mating type, which is prevalent in this species (14) and thus mating may have occurred at some point during the evolution of this strain.

**Table 1:** Genome characteristics of *C. lusitaniae* isolates

| Isolate | CDS | Genes | tRNA | rRNA | Reference |
| --- | --- | --- | --- | --- | --- |
| DSY4606 (P1)<sup>a</sup> | 5676 | 5882 | 197 | 9 | This study |
| DSY4590 (P2) | 5662 | 5869 | 196 | 11 | This study |
| DSY4593 (P3) | 5684 | 5892 | 198 | 10 | This study |
| DSY4661 (P4) | 5679 | 5886 | 197 | 10 | This study |
| DSY4662 (P5) | 5676 | 5883 | 197 | 10 | This study |
| ATCC 42720 | 5936 | 6154 | 217 | NA | (8) |
| CBS 6936 | 5539 | 5740 | 197 | 4 | (11) |

<sup>a</sup>: Isolates designations are both given as laboratory collection and as earlier published in parentheses (7).

A few other characteristics were observed in the genomes of P1 to P5. As mentioned above, a 30 kb region in the Chr 4 in P1, P3 and P4 was duplicated as compared to P2 and P5. This region contains 13 CDS, and among them a putative agglutinin of 2633 amino acids similar to Als2 from *C. albicans*. The significance of this duplication remains unclear. In addition, P1 to P5 isolates contain each 15 separate retrotransposon-like elements divided into 4 different groups, some of which are flanked by nucleotide repeats, referred to as LTRs (Long Terminal Repeats; File S2). So far, the presence of retrotransposons was not firmly reported in this species. Lastly, we observed the expansion of specific protein types in each of these isolates. For example, seven distinct proteins similar to the NADPH-dependent methylglyoxal reductase *GRP2* from *C. albicans* were detected (File S2). *GRP2* in *C. albicans* was shown to be involved in oxidative stress response (15) and is co-regulated with other azole-responsive genes (16). The *C. lusitaniae GRP2*-like genes in P1-P5 exhibit 94-58% similarity with each other. So far, no equivalent of gene family expansion in this type of genes were reported in other *Candida* species.

### Comparative analysis between the P1 to P5 genomes

In order to identify mutations associated with drug resistance in the sequenced *C. lusitaniae* genomes, all 5 genomes were aligned and nucleotide differences in coding and non-coding regions were recorded taking isolate P1 as a reference (see Material and Methods). We showed earlier that isolate P1 was likely to be the parent of all subsequently isolated *C. lusitaniae* samples from the treated patient (isolates P2-P5). Table 2 summarizes the occurrence of nucleotide changes existing between isolates P2 to P5 taking P1 as a genome reference. In general, one can observe a low level of variation between the genomes, even if yeast samples were recovered within a 5 months period. A total of 18 non-synonymous SNPs were observed, among which 13 were unique. We counted 166 SNPs/indel in intergenic sequences between the isolates and 14 other indels in ORFs, some of them (5) causing unique frameshifts and truncations (File S3).

**Table 2:** Occurrence of nucleotide changes in isolates P2-P5 as compared to P1.

|  | non syn SNPs |  |  |  | Intergenic SNPs/indel |  |  |  | ORF Indel/frameshifts |  |  |  |
| --- | --- | --- | --- | --- | --- | --- | --- | --- | --- | --- | --- | --- |
|  | P2 | P3 | P4 | P5 | P2 | P3 | P4 | P5 | P2 | P3 | P4 | P5 |
| **Chr 1** | 1 | 2 | 2 | 1 | 9 | 6 | 6 | 6 | 1 | 0 | 1 | 1 |
| **Chr 2** | 2 | 0 | 1 | 2 | 9 | 5 | 6 | 3 | 0 | 1 | 2 | 1 |
| **Chr 3** | 0 | 3 | 0 | 0 | 10 | 5 | 5 | 6 | 0 | 1 | 0 | 0 |
| **Chr 4** | 0 | 0 | 0 | 0 | 15 | 11 | 8 | 9 | 0 | 1 | 2 | 0 |
| **Chr 5** | 0 | 1 | 0 | 0 | 5 | 4 | 4 | 3 | 0 | 1 | 0 | 0 |
| **Chr 6** | 0 | 0 | 0 | 3 | 2 | 2 | 2 | 1 | 0 | 0 | 1 | 1 |
| **Chr 7** | 0 | 0 | 0 | 0 | 3 | 3 | 2 | 2 | 0 | 0 | 0 | 0 |
| **Chr 8** | 0 | 0 | 0 | 0 | 6 | 4 | 2 | 2 | 0 | 0 | 0 | 0 |
| **Total** | 3 | 6 | 3 | 6 | 59 | 40 | 35 | 32 | 1 | 4 | 6 | 3 |
| **Total all isolates** | 18 |  |  |  | 166 |  |  |  | 14 |  |  |  |

Table 3 includes the 25 different ORFs of isolates P2 to P5 that differ from isolate P1. Two in-frame insertions occurred in two proteins similar to transcription factors (*CZF1*, *SPT4*) in isolates P3 and P5. These insertions could potentially exert positive or negative effects on the function of these factors, which remains to be addressed. Other insertions occurred in isolates P2, P4 and P5 in proteins similar to *BPH1* (a protein playing a role in protein sorting), *BNR1* (cytoskeleton proteins), *CBP1* (corticosteroid-binding protein) and to a protein with unknown function (similar to CLUG_03676 from *C. lusitaniae* ATCC 42720). Six frameshifts mutations occurred in theses genome comparisons. Interestingly, two of these were proteins with potential functions in cell wall biogenesis. The first one was a frameshift mutation in EJF17_2079 (isolate P4) with similarity to *UTR2* (extracellular glycosidase) resulting in a truncation of the 26 amino acids of the full protein. The significance of this alteration is unknown, however the C-terminal end of Utr2p contains a signal sequence important for GPI-anchoring of the protein. Therefore, one can speculate some functional consequence of this protein truncation in isolate P4. The second one was a frameshift mutation in EJF15_20956 and EJF17_20956 from isolates P3 and P4 with similarity to *GAS4* (a 1,3-beta-glucanosyltransferase). This mutation also results in a truncation of the full protein at the C-terminus but of only 3 amino acids. Gas4p contains also a C-terminal signal sequence important for GPI-anchoring of the protein. This short truncation may have limited impact on protein function, however this needs still further verification. The 4 other frameshifts mutation occurred in ORFs with other functions in P2 to P5 isolates, all resulting in protein truncations. It is difficult to predict to which extent these truncations could affect the current phenotypic characterization of the recovered isolates.

**Table 3:** ORFs alterations in isolates P1-P5 as compared to P1.

| P2-P4 altered loci | Isolate | Description | Amino Acid Change | Protein Effect | CDS Codon Number |
| --- | --- | --- | --- | --- | --- |
| EJF18_20133 | P2, P5 | similar to ERG4_YEAST,Delta(24(24(1)))-sterol reductase |  | Truncation | 412 |
| EJF18_60340 | P5 | similar to ERG3_CANAX,Delta(7)-sterol 5(6)-desaturase |  | Truncation | 308 |
| EJF15_10551, EJF17_10551 | P3, P4 | similar to MRR1_CANAL,Multidrug resistance regulator 1 | V -> G | Substitution | 668 |
| EJF15_10714, EJF17_10714, EJF18_10714 | P3, P4, P5, P2 | similar to BSC5_YEAST,Bypass of stop codon protein 5 | F -> L | Substitution | 447 |
| EJF17_20482 | P2 | similar to FKS1_YEAST,1,3-beta-glucan synthase component FKS1 | S -> Y | Substitution | 638 |
| EJF16_20482 | P4 | similar to FKS1_YEAST,1,3-beta-glucan synthase component FKS1 | S -> Y | Substitution | 631 |
| EJF18_20482 | P5 | similar to FKS1_YEAST,1,3-beta-glucan synthase component FKS1 | S -> P | Substitution | 638 |
| EJF15_30508 | P3 | similar to OPT2_YEAST,Oligopeptide transporter 2 | P -> L | Substitution | 895 |
| EJF15_30569 | P3 | similar to MSH2_YEAST,DNA mismatch repair MSH2 | V -> G | Substitution | 186 |
| EJF15_30782 | P3 | similar to 60S ribosomal protein | E -> K | Substitution | 206 |
| EJF15_50669 | P3 | similar to hypothetical protein CLUG_04164 Clavispora lusitaniae ATCC 42720 | P -> S | Substitution | 25 |
| EJF18_60098 | P5 | similar to CBP1_CANAL,Corticosteroid-binding protein | A -> K | Substitution | 531 |
| EJF18_60098 | P5 | similar to CBP1_CANAL,Corticosteroid-binding protein | K -> A | Substitution | 536 |
| EJF18_20361 | P5 | similar to STP4_CANAL,Transcriptional regulator STP4 | QF -> QFQYPGGTKKSQRNQVA GCCRLCQAQF | Insertion | 306 |
| EJF15_30283 | P3 | similar to CZF1_CANAL, Zinc cluster transcription factor CZF1 | N -> NN | Insertion | 150 |
| EJF17_40359 | P4 | similar to BPH1_YEAST,Beige protein 1 | EE -> E | Insertion | 1181 |
| EJF17_40476 | P4 | similar to hypothetical protein CLUG_03676 Clavispora lusitaniae ATCC 42720 | Q -> QQ | Insertion | 141 |
| EJF16_40795 | P2 | similar to BNR1_CANAL,Formin BNR1 | HEAKD -> H | Insertion | 1777 |
| EJF18_60098 | P5 | similar to CBP1_CANAL,Corticosteroid-binding protein | D -> DKEDRD | Insertion | 540 |
| EJF17_10341 | P4 | similar to YD338_YEAST,Uncharacterized transporter YDR338C |  | Frame Shift | 39 |
| EJF18_10886, EJF16_10886 | P5, P2 | similar to predicted protein CLUG_00859 Clavispora lusitaniae ATCC 42720 |  | Frame Shift | 89 |
| EJF17_20798 | P4 | similar to UTR2_CANAL,Extracellular glycosidase UTR2 |  | Frame Shift | 401 |
| EJF15_20956, EJF17_20956 | P3, P4 | similar to GAS4_YEAST,1,3-beta-glucanosyltransferase GAS4 |  | Frame Shift | 175 |
| EJF15_50735 | P3 | similar to LGUL_YEAST,Lactoylglutathione lyase |  | Frame Shift | 268 |
| EJF17_60452 | P4 | similar to APE2_CANAL,Aminopeptidase 2 |  | Frame Shift | 725 |

With regards to single amino acid substitutions detected in the genome comparisons, we first observed changes in four ORFs including EJF15_30508 (similar to *OPT2*, an oligopeptide transporter), EJF15_30569 (similar to *MSH2*, a DNA mismatch repair gene), EJF15_30782 (similar to a 60S ribosomal protein), EJF15_50669 (similar to the hypothetical protein CLUG_04164 from *C. lusitaniae* ATCC 42720) and EJF18_60098 (similar to *CBP1*, a corticosteroid-binding protein). Out of these, the change V186G in EJF15_30569 (similar to *MSH2*) from isolate P3 if of potential interest. Yeast *MSH2* is part of a complex involved in DNA repair and *MSH2* mutations have been shown to contribute to increasing mutation rates in several fungal species (17, 18). The V186G substitution in EJF15_30569 lies near a domain referred to as the connector domain that is important for the function of Msh2p (19). It is therefore possible that the V186G substitution from isolate P3 affects Msh2p function and consequently the frequency at which mutations can occur in isolate P3 as compared to P1.

While inspecting these data in the perspective of antifungal drug resistance, mutations in several genes associated with the specific drug resistance profiles of P2 to P5 isolates could be highlighted and are summarized below:

i) V668G in EJF15_10551 and EJF17_10551 from isolates P3 and P4: these two genes encode a protein similar to MRR1_CANAL (Multidrug resistance regulator 1 of *C. albicans*). In *C. albicans*, this protein is known as a regulator of *MDR1*, a Major Facilitator efflux transporter that mediates fluconazole resistance (20). Gain of function (GOF) mutations in *MRR1* result in strong upregulation of this transporter and, consequently, azole resistance. *MFS7* (homolog of *MDR1* in *C. lusitaniae*) was previously reported to be upregulated in isolates P3 and P4 as compared to P1 (7). We therefore propose that the V668G substitution in EJF15_10551 and EJF17_10551 (now referred as to *MRR1*) is a GOF mutation that mediates *MFS7* upregulation in P3 and P4.

ii) S638Y, S631Y and S638P in EJF17_20482, EJF16_20482 and EJF18_20482 from P2, P4 and P5: these genes encode proteins similar to FKS1_YEAST (1,3-beta-glucan synthase). Fks1p is a β-1,3 glucan synthase, a critical enzyme in the biosynthesis of β-1,3 glucans in fungi. It is also the target of candins (21). We reported earlier the presence of these distinct mutations in *C. lusitaniae* isolates P2, P4 and P5 and showed that they were causing candin resistance but using *Saccharomyces cerevisiae* as a surrogate system (7). The current sequencing data confirm these different mutations in EJF17_20482, EJF16_20482 and EJF18_20482 (now referred as to *FKS1*) that were previously deduced from specific Sanger sequencing reactions.

iii) Truncations in EJF18_20133 and EJF18_60340 from isolates P2 and P5: these genes encode proteins similar to ERG4_YEAST (Delta(24(24(1)))-sterol reductase) and ERG3_CANAX (Delta(7)-sterol 5(6)-desaturase), respectively. In isolate P2, EJF18_20133 (further referred as to *ERG4*) underwent a nucleotide change (C to A) converting the TCG codon (Ser^412^) into a stop codon (TAG: *erg4^amber^*), thus truncating the full Erg4p by 49 amino acids. While the same mutation was detected in isolate P5, another distinct mutation in EJF18_60340 (now referred as to *ERG3*) was revealed in the same isolate. *ERG3* underwent a nucleotide change (C to T) converting the CAA codon (Gln^308^) into a stop codon (TAA: *erg3^ochre^*), thus truncating the full Erg3p by 59 amino acids. Both P2 and P5 were reported as resistant to AmB (7). Thus, both *ERG4* and *ERG3* truncations may have resulted in altered sterol composition in these isolates and eventually leading to the depletion of ergosterol, which is a known factor mediating AmB resistance (22).

Taking these comparative descriptions together, we summarized the potential associations existing between the isolate genotypes and the reported antifungal drug resistance profiles in Fig. 2 considering only coding sequences for comparisons. Isolate P1 was previously shown as the earliest yeast recovered from the patient at early stages of antifungal treatment and isolates P2 to P5 were timely sequential isolates. Given the high nucleotide resemblance between the isolates and given that all isolates exhibit the same SNP as compared to P1 (*BSC5* F447L), the data confirm that P1 is the ancestor of P2 to P5. However, it is less likely that each isolate represents a progeny from the other. This conclusion is supported by the different SNP profiles found in the isolates (Table 3). Interestingly, P3 and P4 share the same substitution in *MRR1* (V668G) and the same frameshift mutation in EJF15_20956 and EJF17_20956 (*GAS4*), thus pointing to a common ancestor. P2 and P5 also share SNPs leading to the same truncation in *ERG4* (S412*) and the same frameshift mutation in EJF14_10886, which suggests that both isolates originated from a common ancestor. However, each of these isolates also contains specific other SNPs, indicating that they evolved in individual directions. To summarize, besides the already known reported *FKS1* mutations, we identified here 3 novel mutations including V668G in *MRR1* possibly mediating FLC resistance, and S412* in *ERG4* and Q308* in *ERG3* that could be responsible for AmB resistance.

**Figure 2:**
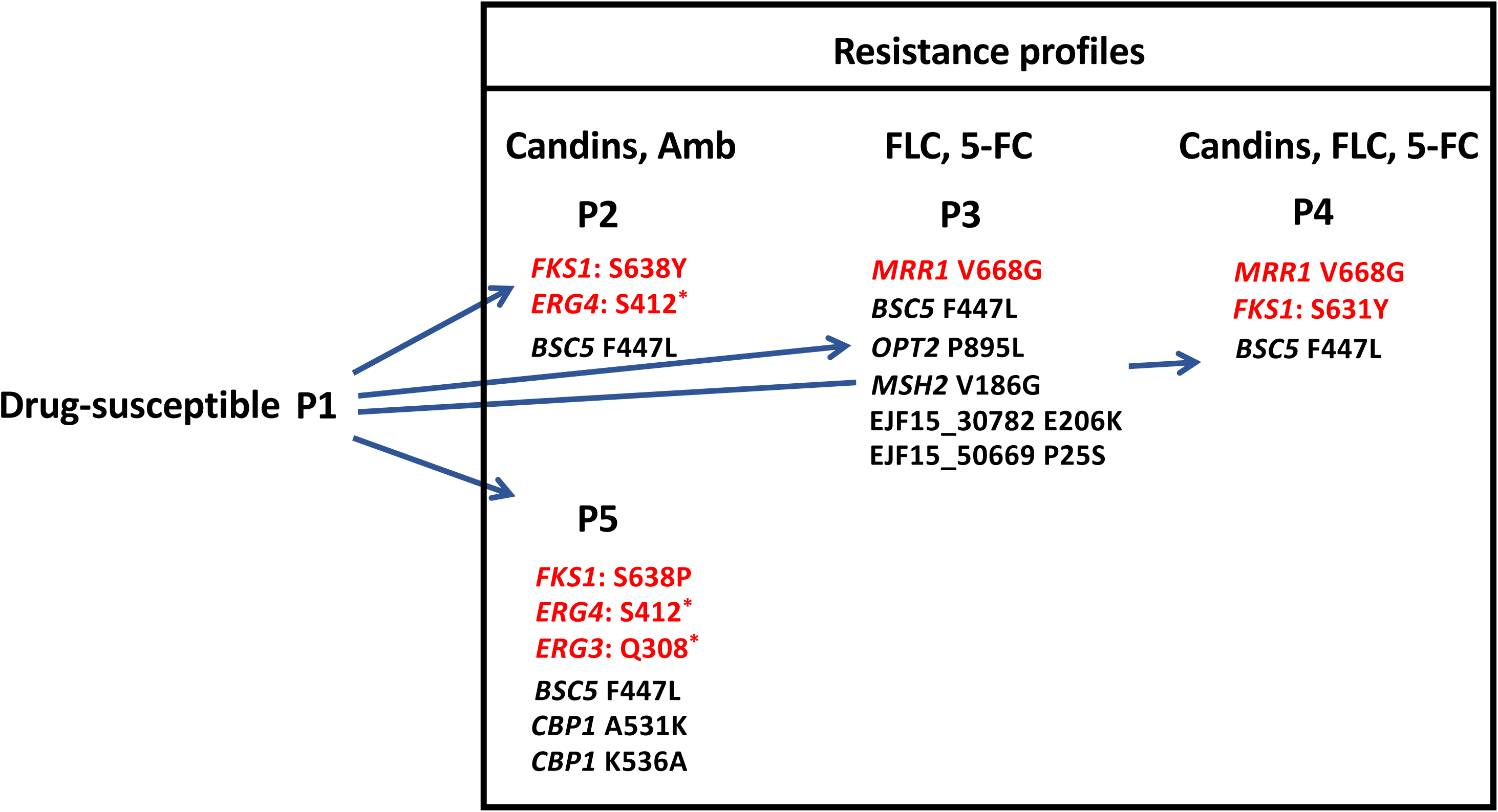
Relationships between isolates drug resistance phenotypes and genotypes. Only SNP variants in coding regions are reported here. Red labels indicate SNPs that may cause drug resistance. S412* and Q308* signify change of serine and glutamine into a stop codon.

### Role of *MRR1* in FLC resistance

Our data suggested that *MRR1* (EJF14_10551) could be involved in FLC resistance in *C. lusitaniae* due to the occurrence of a V668G substitution. To address this question, we first undertook a RNAseq analysis in isolates P3 and P4 grown under normal conditions and compared the data with P1 (see Material and Methods). Table 4 lists the genes up- and down-regulated in P3/P4 as compared to P1. According to our selection (1.5-fold expression change as compared to P1), there were 54 and 30 genes up- and down-regulated, respectively, in P4. When comparing P3 with P1, 16 genes were inversely regulated keeping this threshold. Using a Gene Ontology (GO)-terms enrichment analysis based on gene annotations of *C. albicans* homologs, significant enrichment of functions related to azole transport and oxido-reductase activity were detected among commonly upregulated genes in P3 and P4 (File S4). Highly upregulated genes (215- to 57-fold in P4 and P3 vs P1) involved in these biological processes include alcohol dehydrogenases (EJF14_60130, EJF14_40004 similar to *ADH6* and *ADH7*) and a NADPH-dependent methylglyoxal reductase (EJF14_20072 similar to *GRP2*). *MFS7* was among these highly upregulated genes (17- and 15-fold upregulation in P4 and P3 vs P1) in isolates containing the *MRR1* V668G substitution. *MRR1* itself was significantly upregulated in P3/P4 isolates (>1.5-fold vs P1). These expression profiles are consistent with the imprint of *MRR1* GOF mutations in *C. albicans* on the transcriptome of this species (20). We therefore suggest that the *C. lusitaniae MRR1* V668G substitution is a GOF mutation which impacts on the expression of different target genes, and among them *MFS7*.

**Table 4:**
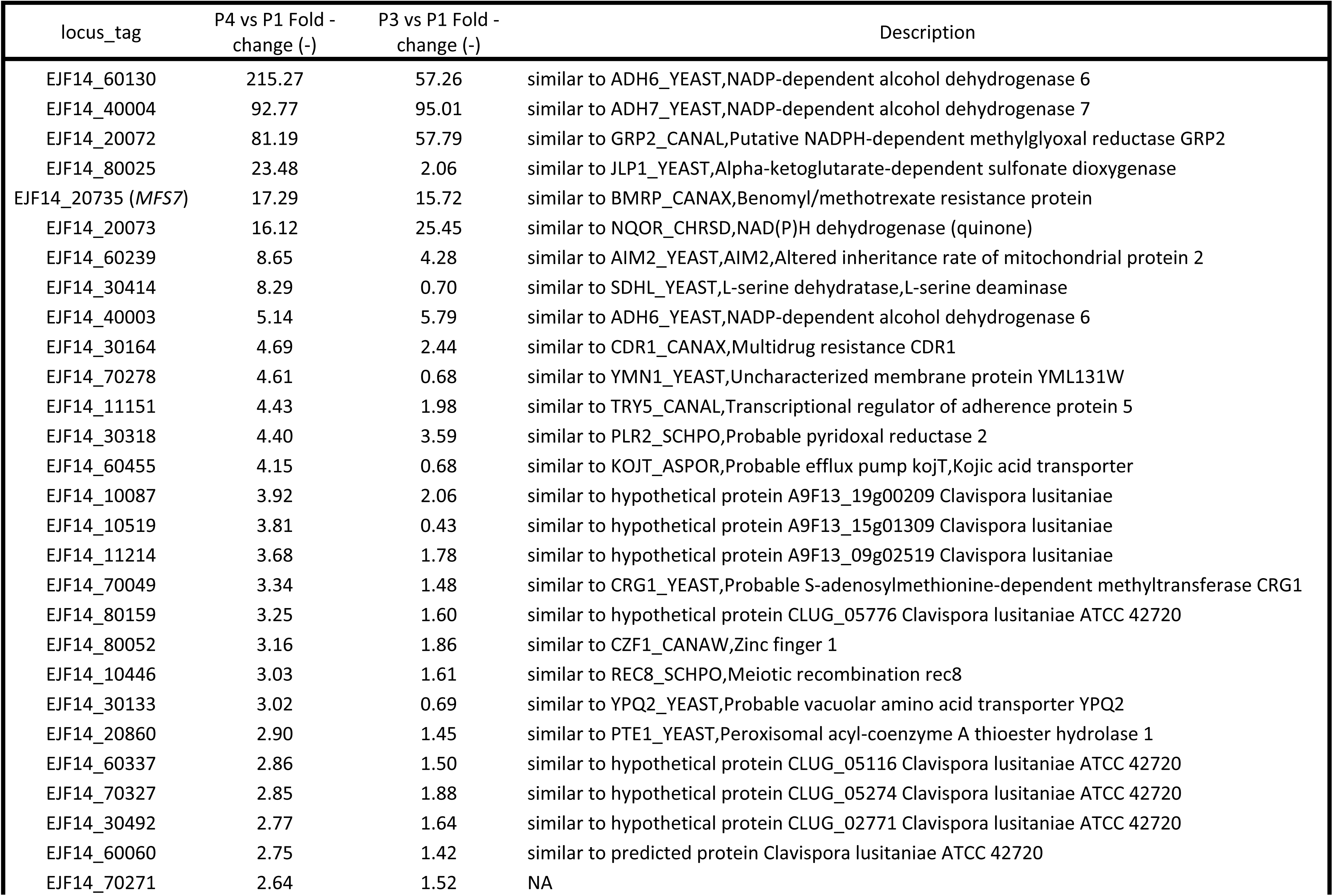
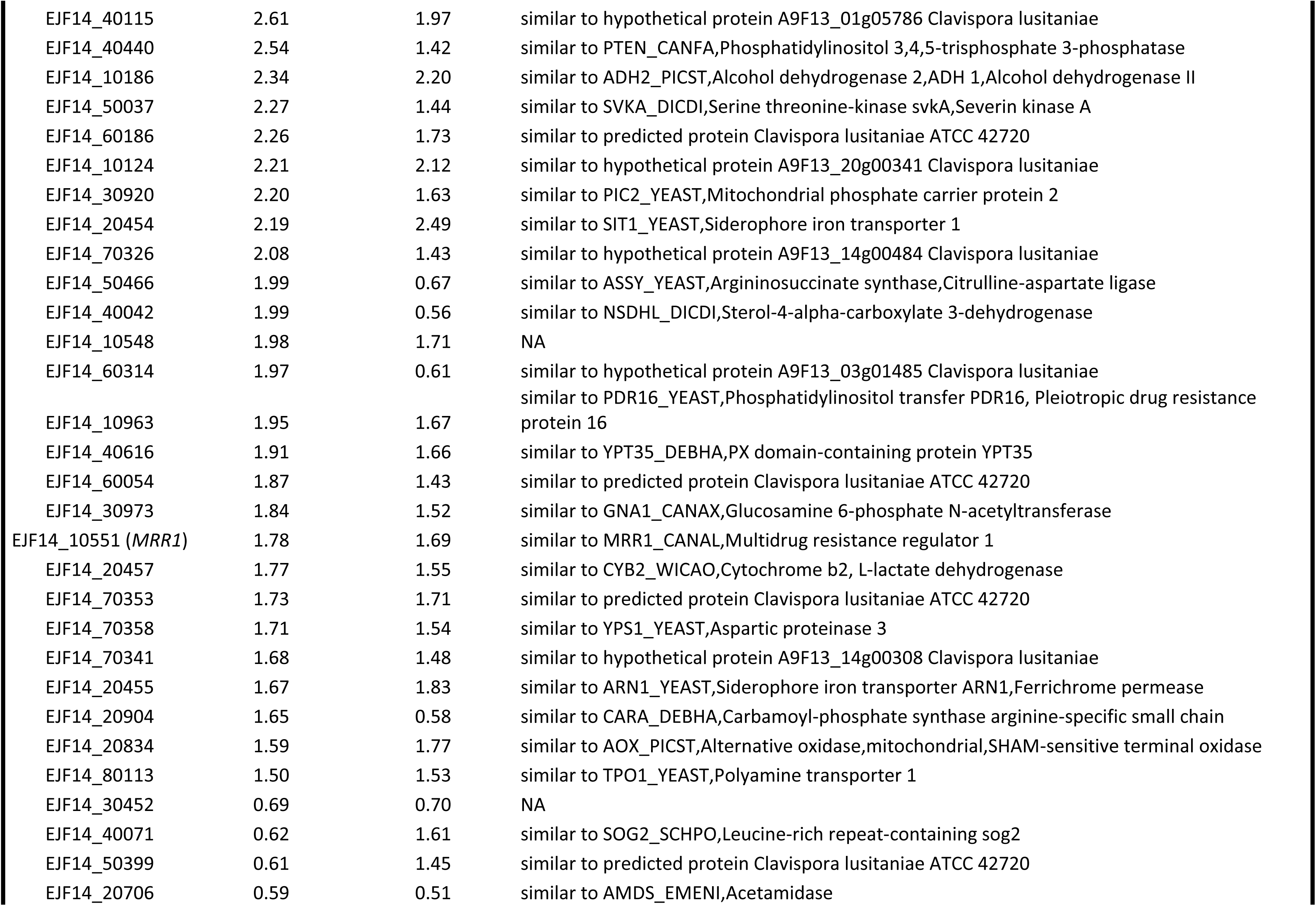
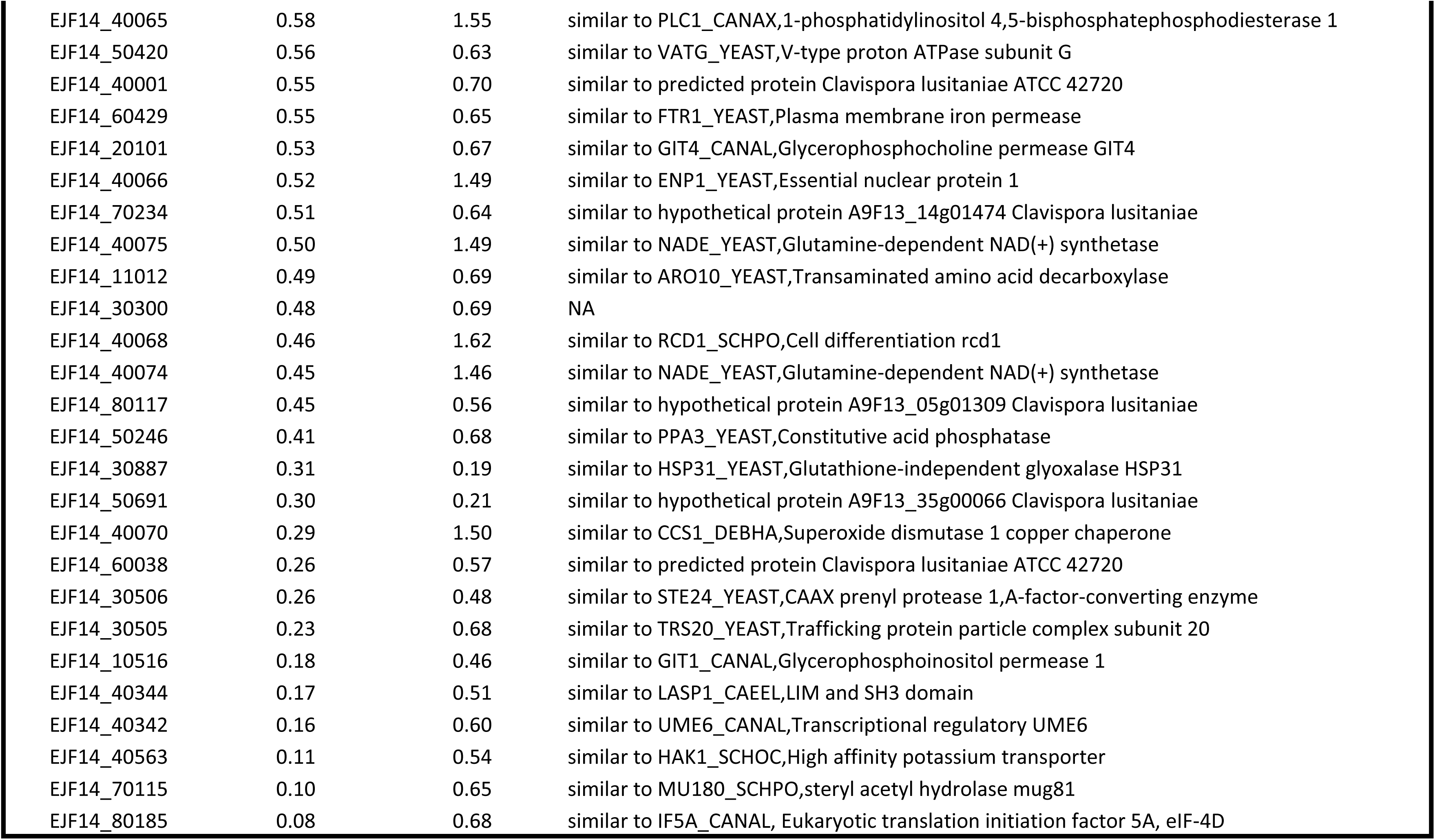
Transcriptomic data of azole-resistant isolates P3 and P4 as compared to the susceptible isolate P1.

In order to further address the role of *MRR1* and *MFS7* in the drug resistance profiles, we first attempted gene knock-out approaches. Preliminary results using homologous recombination with a dominant marker (*SAT1*) flanked by *MFS7-* and *MRR1* 5’- and 3’-regions proved as inefficient. We therefore used a CRISPR-Cas9 approach with a guide RNA specific for *MFS7* and *MRR1* using *in vitro* reconstituted RNA-protein complexes (RNPs) and repair fragments with the dominant marker by *NAT1* flanked by *MFS7-* and *MRR1* 5’- and 3’-regions (see Material and Methods). CRISPR-Cas9 and corresponding repair fragments were used in isolates P1 and P3 and expected mutants were verified as described (see Materials and Methods). Fig. 3A shows that the deletion of both *MFS7* and *MRR1* in the background of azole-susceptible isolate P1 did not significantly alter FLC susceptibility assessed both by serial dilution assays and MIC measurements (MIC values fluctuated between 1- and 0.25 µg/mL FLC for all P1 and P1-derived mutants; see File S5 for full MIC data). On the contrary, deletion of both *MFS7* and *MRR1* in the background of the azole-resistant isolate P3 significantly altered azole susceptibility: while the initial isolate P3 exhibited a MIC value of 64 µg/mL FLC, deletion of *MRR1* resulted in 64-fold MIC decrease (MIC = 1 µg/mL FLC; File S5) and deletion of *MFS7* in a 8-fold MIC decrease (MIC = 8 µg/mL FLC; File S5). We addressed the effect of *MRR1* deletion on *MFS7* expression by qPCR and, while *MFS7* expression was increased by 28-fold between the isolates P1 and P3, it was decreased by 14-fold in the absence of *MRR1* between P3 and its *MRR1*-derived mutant (Fig. 4A). These findings indicate that the *MRR1* allele of P3 containing the V668G substitution mediates *MFS7* regulation in *C. lusitaniae*.

**Figure 3:**
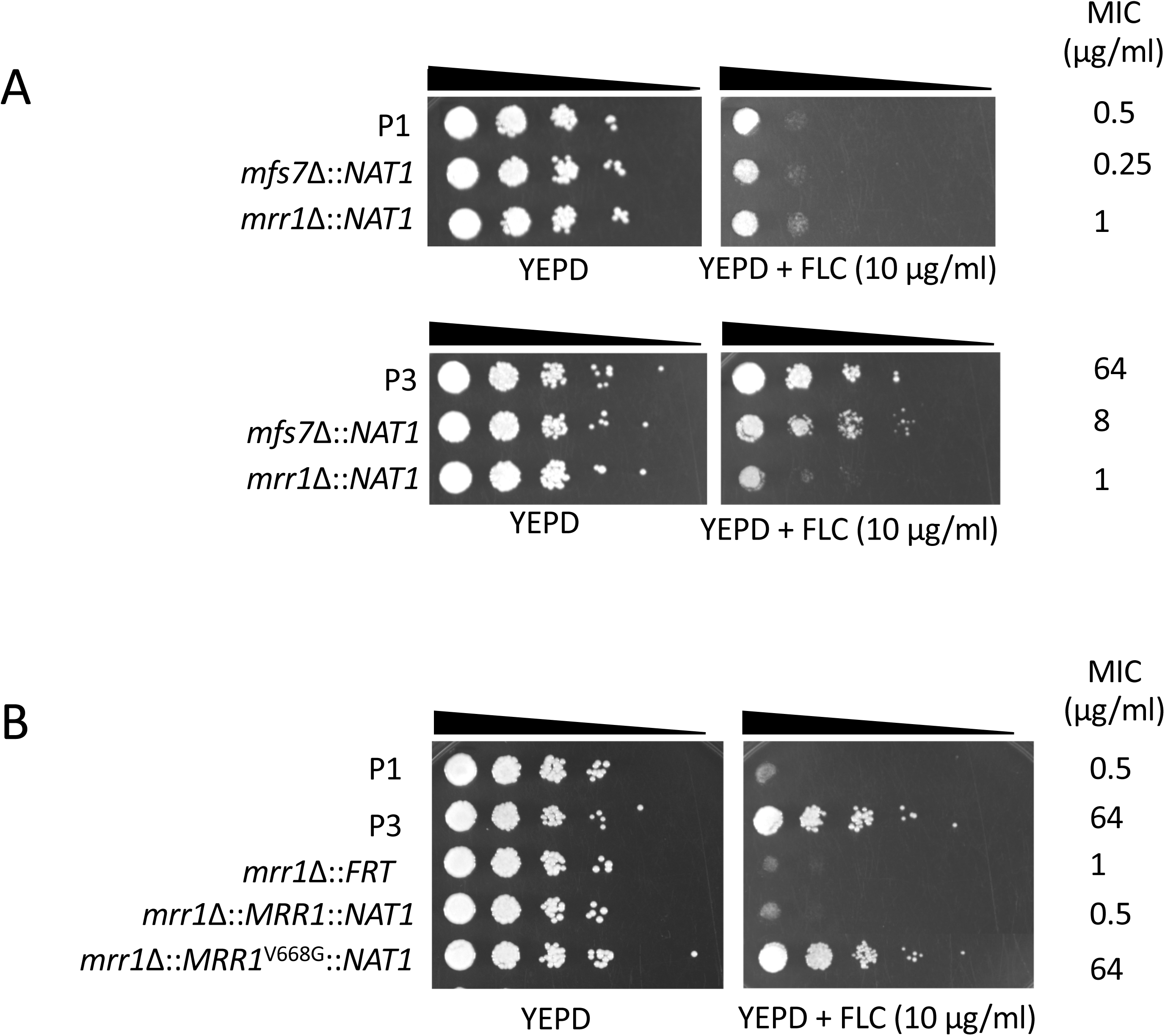
*MRR1* and *MFS7* mediate FLC resistance in *C. lusitaniae*. **Panel A:** Ten-fold serial dilutions were performed starting with an inoculum of about 10^5^ cells. Mutants for *MRR1* and *MFS7* in P3 correspond to DSY5240 and DSY5242 and in P1 to isolates DSY5246 and DSY5248, respectively. MIC values were obtained by MIC measurements using the SYO system as described in Material and Methods. **Panel B:** Reversion of *MRR1* deletion. Mutants for *MRR1* correspond to DSY5416. Revertants for *MRR1* wild type allele and *MRR1* GOF allele (*MRR1*^V668G^) correspond to isolates DSY5437 and DSY5438, respectively.

**Figure 4:**
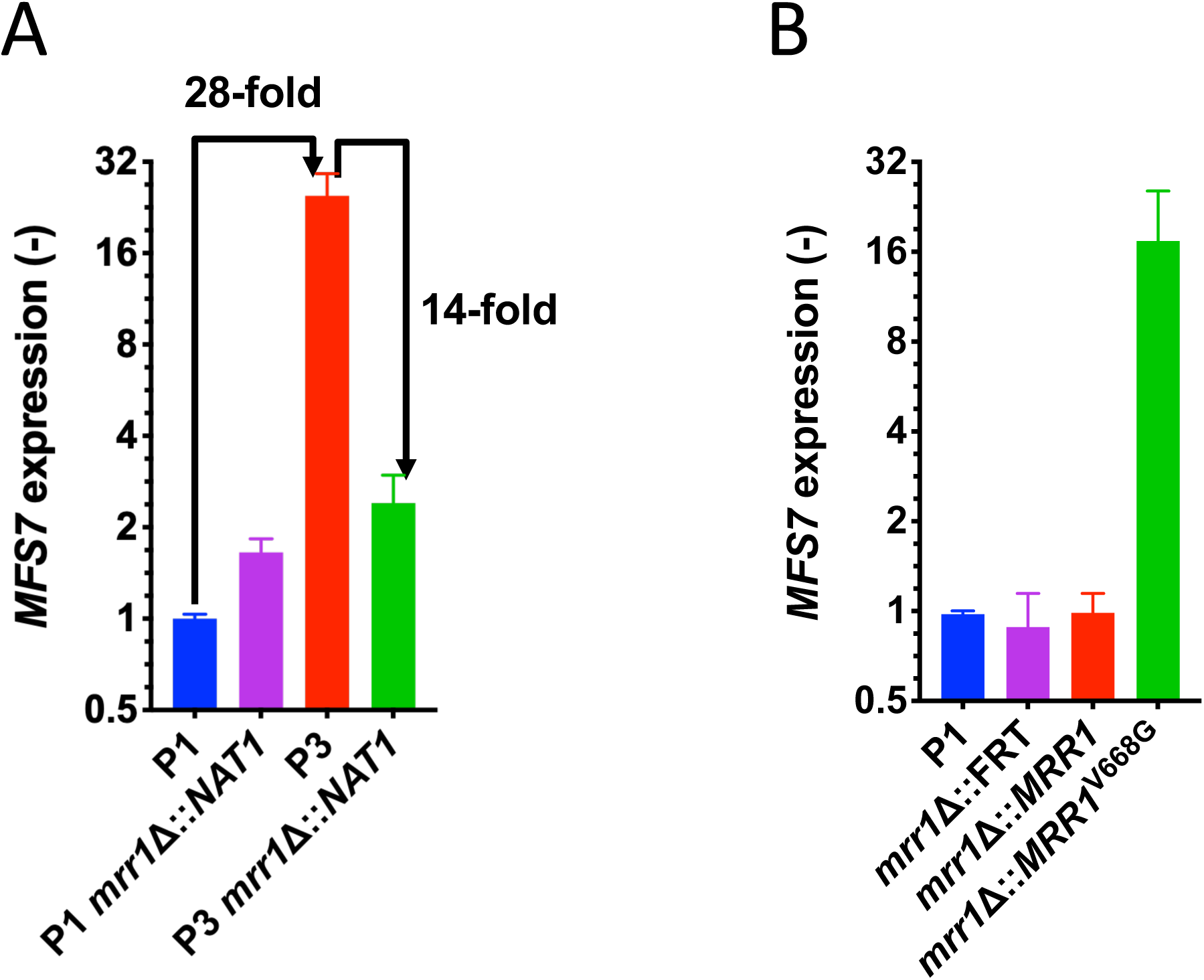
Expression of *MFS7* in *C. lusitaniae.* **Panel A:** *MFS7* expression in P1 and P3 and derived *MRR1* mutants (DSY5246 and DSY5240). *MFS7* expression fold-changes are indicated between specific isolates. **Panel B:** *MFS7* expression in *MRR1* revertants (DSY5437: *mrr1*Δ::*MRR1* and DSY5438: *mrr1*Δ::*MRR1*^V668G^) derived from DSY5416 (*mrr1*Δ::FRT). *MFS7* expression was calculated relative to the initial isolate P1. qPCR were performed with biological triplicates as described in Materials and Methods.

To demonstrate a genetic link between the presence of a *MRR1* GOF mutation and azole resistance, we next constructed revertants from a *MRR1* mutant in which the *MRR1* alleles from P1 (wild type allele) and P3 (GOF allele) were re-introduced. This experiment required the recycling of the *NAT1* dominant marker, thus a *MAL2*-dependent flipper system permitting excision of the *NAT1* marker was designed and a separate *MRR1* mutant lacking the selection maker was produced (DSY5416, see Material and Methods). CRISPR-Cas9 combining two guide RNAs targeting flanking *MRR1* regions were used with *MRR1* wild type and GOF alleles on the *NAT1* dominant marker to produce final revertants. Fig. 3B shows that the presence of the *MRR1* GOF allele alone could restore azole resistance (MIC = 64 µg/mL FLC; File S5) and thus demonstrates the association between the *MRR1* GOF mutation V668G and azole resistance. Consistently, high *MFS7* expression was restored from the *MRR1* deletion mutant in the presence of the *MRR1* GOF mutation and not with the *MRR1* wild type allele (Fig. 4B).

### Role of *MRR1* in 5-FC resistance

We showed earlier in the *C. lusitaniae* isolates recovered from a treated patient that FLC resistance was associated with 5-FC resistance, even if the patient was not exposed to the pyrimidine analog. 5-FC resistance in *C. lusitaniae* is mediated by mutations occurring in genes involved in 5-FC transport and metabolism (*FCY1*, *FCY2*) (23). In other species such as *C. albicans*, other genes (*FUR1* encoding uracil phosphoribosyltransferase) contains mutations resulting in 5-FC resistance (24). None of these genes contained mutations in P2 to P5 isolates. This raised the hypothesis that 5-FC resistance could be mediated by alternative mechanisms, and especially by *MFS7* upregulation, since it results in FLC resistance that itself correlates with 5-FC resistance. We tested this hypothesis by first expressing *MFS7* in a heterologous system in which major *C. albicans* multidrug transporters (*CDR1*, *CDR2*, *MDR1*, *FLU1*) were inactivated (25). We observed by serial dilution spotting assays (Fig. 5A) that FLC resistance occurred when *MFS7* was overexpressed as did the *C. albicans* ABC-transporter *CDR1*, as expected. Interestingly, *MFS7* overexpression resulted in a specific 5-FC resistance since it was not observed by *CDR1* overexpression (Fig. 5A). We also confirmed by GFP tagging that *MFS7* localized principally to cell structures in *C. albicans* believed as cell membranes (Fig. 5B). These data suggest *MFS7* as a cell membrane-associated transporter for 5-FC. We took advantage of *MRR1* and *MFS7* mutants to address their susceptibility to 5-FC. Fig. 6A shows that deletion of both genes resulted in a 64-fold decrease of MIC as compared to the parent strain P3. In addition, the reversion of the *MRR1* deletion by a *MRR1* GOF allele restored 5-FC resistance (Fig. 6B; File S5). Taken together, our data indicate for the first time that *MRR1* is responsible for 5-FC resistance in *C. lusitaniae* and that this resistance is mediated by *MFS7*, especially when upregulated. This novel mechanism is likely to explain the cross-resistance between the two drug classes in *C. lusitaniae*.

**Figure 5:**
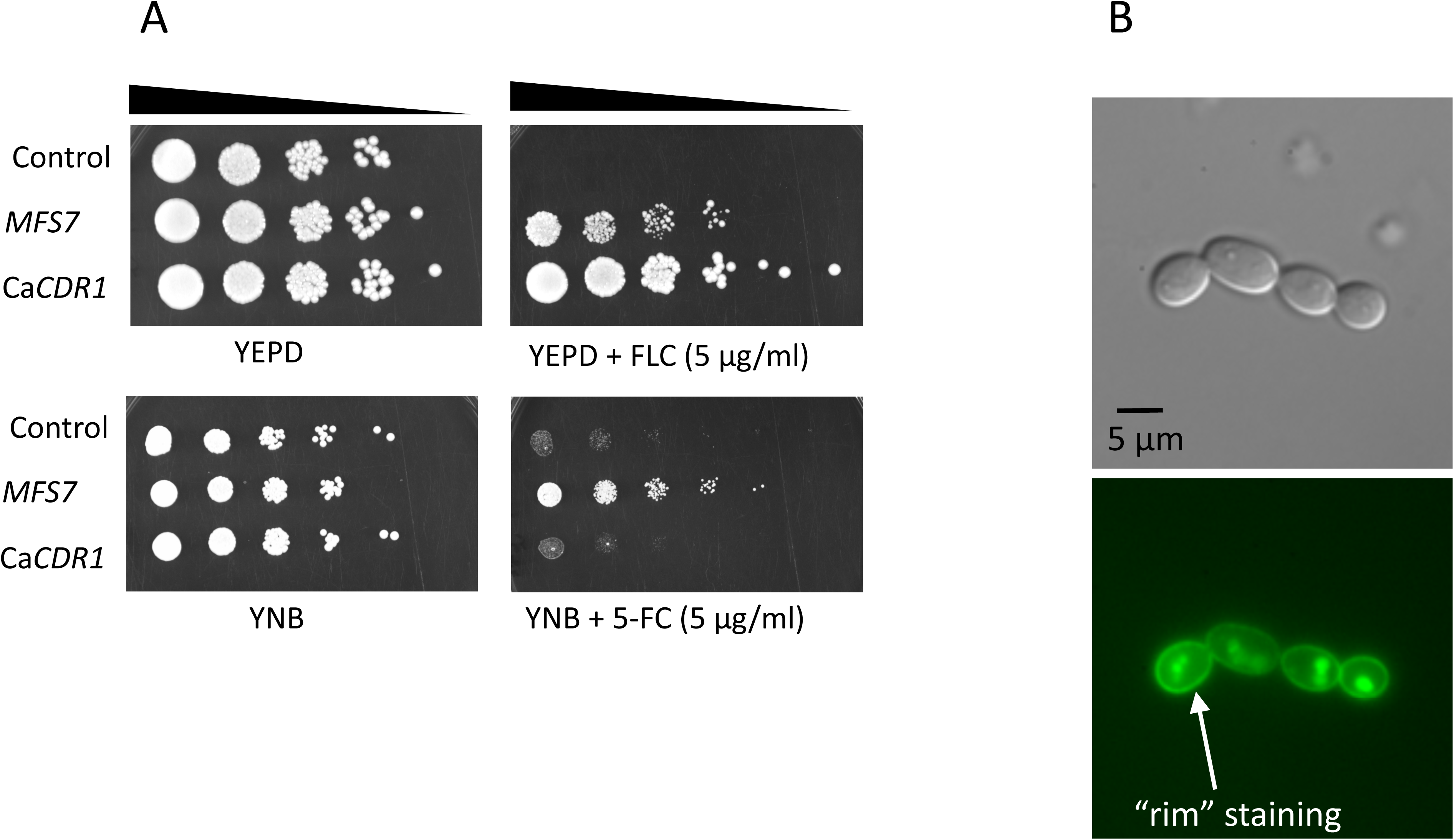
*MFS7* mediates 5-FC resistance. **Panel A**: Ten-fold serial dilutions were performed starting with an inoculum of about 10^5^ cells at the indicated 5-FC concentration. *MFS7* overexpression was obtained by a *CDR1*-MFS7 chimeric construct expressed in DSY5170. Control strain is DSY5169 (See Table S1); *CDR1* overexpression stain is ANY-MDR1-GFP (Ca *CDR1* in Figure caption). **Panel B**: localisation of MFS7-GFP in *C. albicans*. Microscopy was performed with under normal light with differential interference contrast (DIC) microscopy and with epifluorescence as described in Materials and Methods. GFP localization shows the typical “rim” staining of membrane proteins.

**Figure 6:**
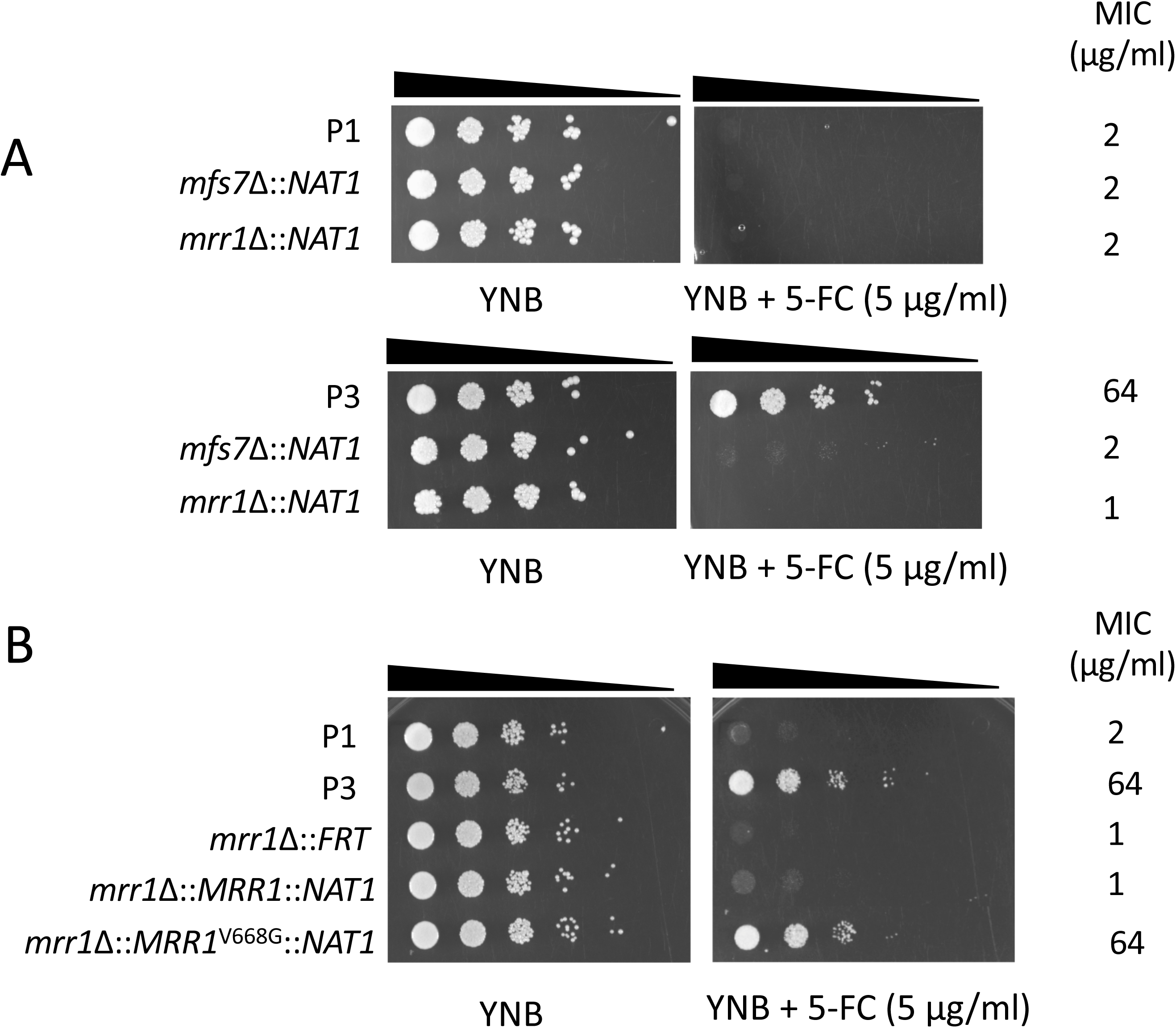
*MRR1* and *MFS7* mediate 5-FC resistance in *C. lusitaniae*. **Panel A:** Ten-fold serial dilutions were performed starting with an inoculum of about 10^5^ cells. Mutants for *MRR1* and *MFS7* in P3 correspond to DSY5240 and DSY5246 and in P1 to isolates DSY5242 and DSY5248, respectively. MIC values were obtained by MIC measurements using the SYO system as described in Material and Methods. **Panel B:** Reversion of *MRR1* deletion. Mutants for *MRR1* correspond to DSY5416. Revertants for *MRR1* wild type allele and *MRR1* GOF allele (*MRR1*^V668G^) correspond to isolates DSY5437 and DSY5438, respectively.

### *ERG3* and *ERG4* loss of functions and AmB resistance of isolates P2 and P5

In our previous report, the mechanisms behind AmB resistance of isolates P2 and P5 remained puzzling. P2 and P5 isolates exhibited Amb MIC of 1 and 2 µg/mL as compared to P1 (MIC = 0.25 µg/mL; File S5). Here we showed by genome comparisons that the two isolates contained SNPs converting codons of *ERG3* and *ERG4* into stop codons. P2 only exhibited a truncation in Erg4p while P5 carried truncations in both Erg4p and Erg3p. These mutations could result in loss of functions and therefore interruption of ergosterol biosynthesis. Fig. 7 shows the sterol biosynthesis pathway and highlights the position of *ERG3* and *ERG4* as well as interference of mutations in P2 and P5 in this pathway. A single *ERG4* loss of function mutation is expected to produce the accumulation of ergosta-5,7,22,24-(28)-tetraenol together with other sterol precursors. The combined *ERG3* and *ERG4* loss of function mutations are expected to yield ergosta-7,22,24-(28)-trienol as major by-product. These expectations could be verified by mass spectrometry analysis of the sterol fractions of P2 and P5. As shown in Table 5, P2 accumulated ergosta-5,7,22,24-(28)-tetraenol up to 98% in the sterol fraction, while P5 accumulated ergosta-7,22,24-(28)-trienol up to 82% of total sterols with other precursors (fecosterol and episterol) resulting from reactions upstream of *ERG3*. The other Amb-susceptible isolates (P1, P3 and P4) exhibited ergosterol as major sterol component (95-98% of total sterols). Taken together, this sterol analysis suggests that the identified *ERG3* and *ERG4* mutations result in loss of functions of the gene products. The accumulation of 2 separate mutations in P5 yields the expected sterol composition. We believe that the absence of ergosterol in P2 and P5 results in their Amb resistance as reported earlier (7). To further address the relationships between phenotypes and genotypes in P2 and P5, *ERG3* and *ERG4* wild type copies were re-introduced in these isolates by a CRISPR-Cas9 approach. Isolate P5 necessitated the restoration of two different wild type gene copies which could be carried out by the use of two different positive selection markers (*NAT1* and *CaHygB*). Ergosterol biosynthesis was restored in P2/P5 revertants with *ERG3* and *ERG4* wild type copies (DSY5441 and DSY5452 in Table 5). As shown in Fig. 8, Amb resistance of P2/P5 was reversed to MIC levels of the initial P1 isolate when *ERG3* and *ERG4* wild type copies were restored in these isolates (MIC= 0.125 and 0.25 µg/mL, File S5). Interestingly, the *ERG4* revertant DSY5444 with the *erg3^ochre^* defective allele exhibits high AmB MIC (1 µg/mL), which is consistent with absence of ergosterol in this isolate and formation of ergosta-7,22-dienol (Table 5), a known sterol metabolite identified in *erg3 Candida* spp mutants (26). The constructed *ERG4* and *ERG3* revertants, while they exhibited changes in Amb MIC values, did not show altered MIC values for other antifungal agents (File S5). This highlights the specificity of the genetic changes and also demonstrates the genetic basis of Amb resistance in isolates P2 and P5.

**Figure 7:**
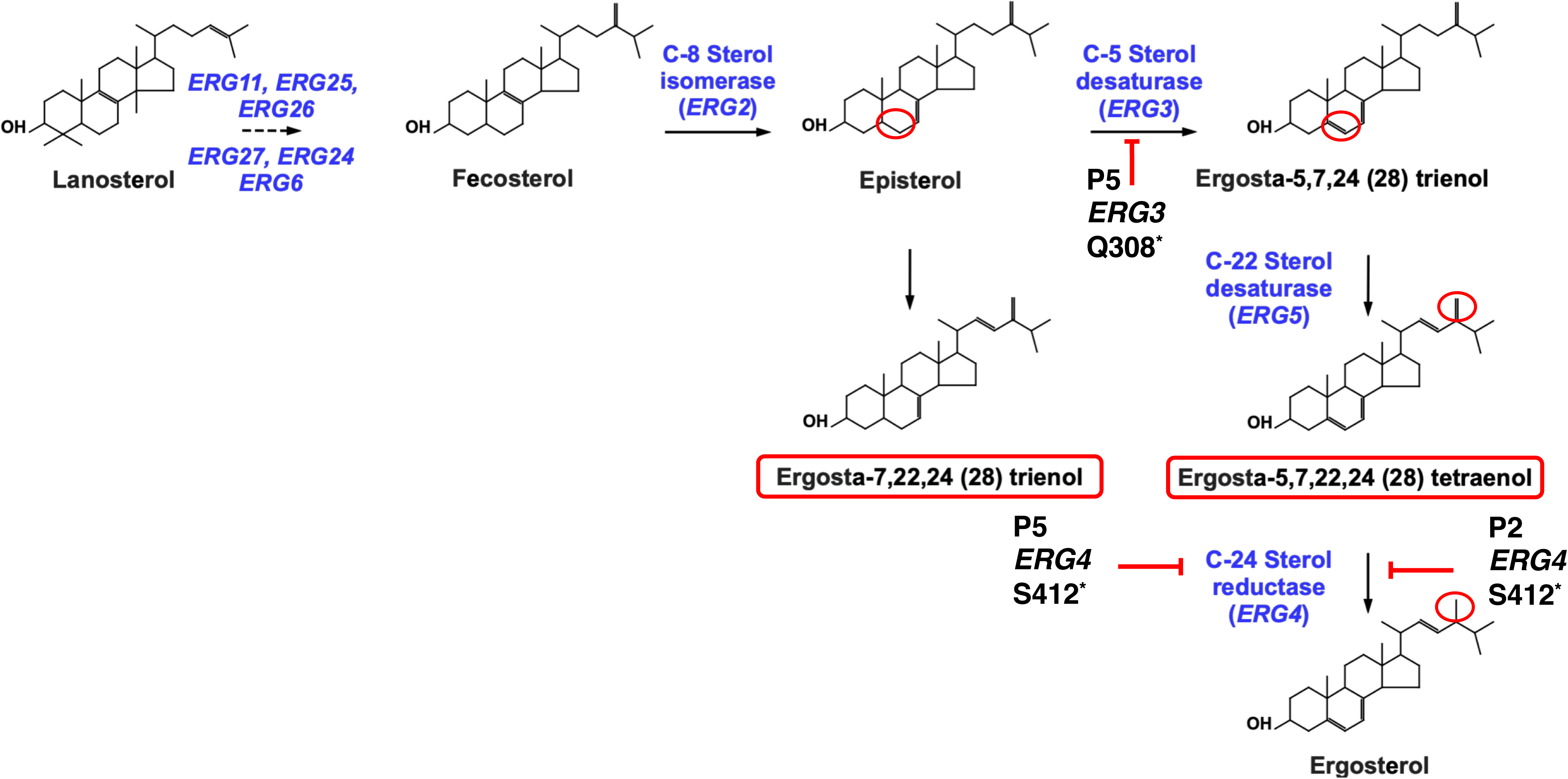
Sterol biosynthesis pathway and defects of isolates P2 and P5. The pathway includes the steps from the substrate lanosterol up to the formation of ergosterol (22). Major genes and sterol intermediates are indicated. *ERG3* and *ERG4* steps are highlighted and red circles show the activity on the sterol molecule (saturation/desaturation). *ERG4* defect in P2 and combined defects in *ERG3* and *ERG4* are indicated with corresponding intermediates accumulations (red rectangles).

**Figure 8:**
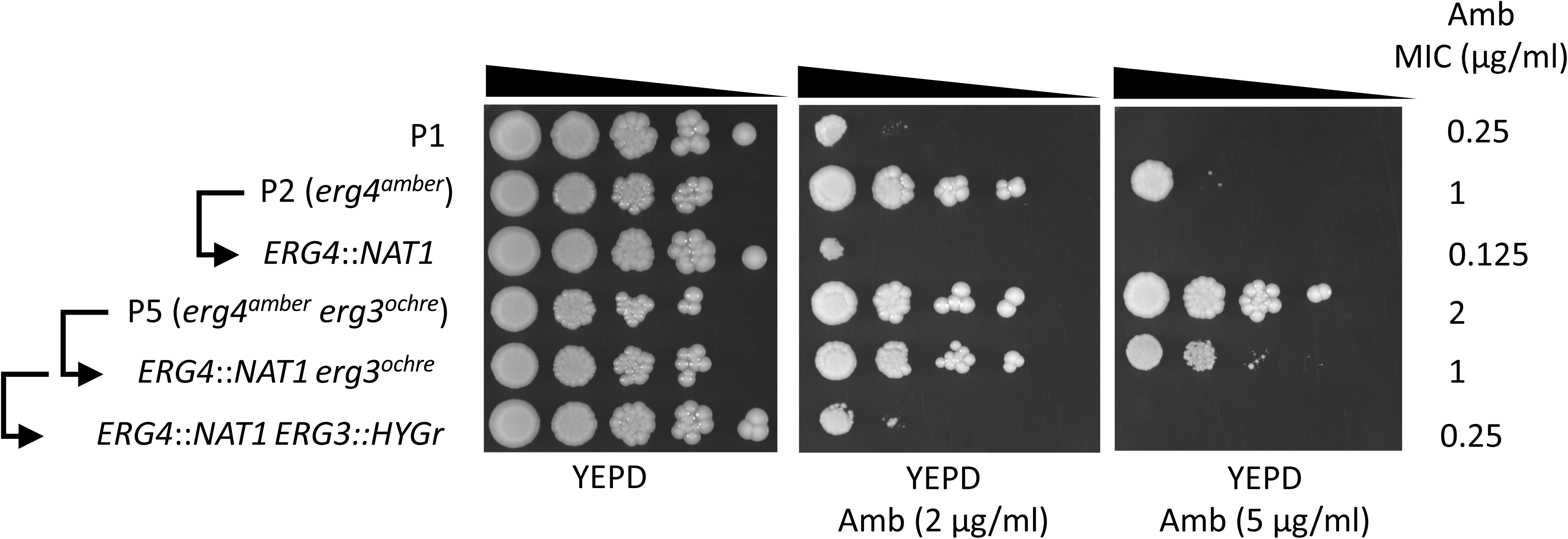
*ERG3* and *ERG4* reversions in isolates P2 and P5. Ten-fold serial dilutions were performed starting with an inoculum of about 10^5^ cells. MIC values were obtained by MIC measurements using the SYO system as described in Material and Methods. The *ERG4* revertants from P2 and P5 are DSY5441 (*erg4^amber^*::*ERG4*::*NAT1*) and DSY5444 (*erg4^amber^*::*ERG4*::*NAT1 erg3^ochre^*). The *ERG4* and *ERG3* revertant from P5 is DSY5452 (*erg4^amber^*::*ERG4*::*NAT1 erg3^ochre^*::*ERG3*::*CaHygB*). Arrows indicate parental relationships between isolates.

**Table 5:** Sterols composition of isolates P1 to P5 and derived mutants

| Sterol metabolite <sup>a</sup> | Sterol content of isolates ( % mean ± S.D) | | | | | | | |
| --- | --- | --- | --- | --- | --- | --- | --- | --- |
|  | P1 | P2<br>*erg4<sup>amber</sup>* | P3 | P4 | P5<br>*erg4<sup>amber</sup>*<br>*erg3<sup>ochre</sup>* | DSY5441<sup>b</sup><br>*ERG4* | DSY5444<sup>b</sup><br>*ERG4*<br>*erg3<sup>ochre</sup>* | DSY5452<sup>b</sup><br>*ERG4 ERG3* |
| Ergosta-8,22-dienol | | | | | | | 5.8 (± 0.2) | |
| Ergosta-5,8,22,24(28)-tetraenol | 0.8 (± 0) | | 0.8 (± 0.1) | 0.7 (± 0.1) | | 0.3 (± 0.0) | | 0.9 (± 0.3) |
| Ergosta-5,8,22-trienol | 0.8 (± 0) | | 0.7 (± 0) | 0.6 (± 0) | | 0.8 (± 0.0) | | |
| Zymosterol | 2.2 (± 0.4) | | 2 (± 0.8) | 2.1 (± 0.1) | | 1.0 (± 0.2) | | |
| **Ergosterol** | **95 (± 0.5)** | | **95.8 (± 0.5)** | **94.8 (± 0.3)** | | **91.3 (± 1.5)** | 0.3 (± 0.1) | **89.2 (± 0.5)** |
| **Ergosta-5,7,22,24(28)-tetraenol** | | **98.5 (± 0.8)** | 0 (± 0.1) | | | 5.5 (± 0.9) | | 9.9 (± 0.2) |
| Ergosta-8,22,24(28)-trienol | | | | | 8.4 (± 0.1) | | | |
| **Ergosta-7,22-dienol** |  |  |  |  |  |  | **65.8 (± 3.1)** |  |
| Fecosterol | 0.1 (± 0) | | 0.3 (± 0.2) | 0.8 (± 0.1) | 0.8 (± 0.4) | | 2.8 (± 0.6) | |
| **Ergosta-7,22,24(28)-trienol** | | | | | **82.3 (± 0.7)** | 0.6 (± 0.2) | 11.9 (± 1.1) | |
| Ergosta-5,7-dienol | 0.7 (± 0.6) | | | 0.3 (± 0.1) | | | | |
| Episterol | | | 0.1 (± 0.2) | 0.3 (± 0.2) | 7.5 (± 1.1) | | 7.1 (± 0.9) | |
| Ergosta-7-enol | | | | | | | 1.3 (± 0.1) | |
| Lanosterol /Obtusifoliol | 0.4 (± 0.2) | | 0.2 (± 0.2) | 0.4 (± 0.2) | | 0.2 (± 0.0) | | |
| 4,4-dimethyl cholesta 8,24 dienol | | | | | | 0.3 (± 0.1) | | |
| Unknown | 0.1 (± 0.1) | 1.5 (± 1) | 0.1 (± 0.1) | | 1.1 (± 0.1) | | 2.2 (± 0.2) | |
| Total | 100 | 100 | 100 | 100 | 100 | 100 | 97.1 | 100 |

<sup>a</sup>: Principal sterols are boldface typed<sup>b</sup>: complete genotypes can be found in Table S2

## Discussion

In this work we showed that the use of genome mining approaches were helpful in the resolution of drug resistance mechanisms in *C. lusitaniae*. Isolates P1 to P5 were recovered within a 5-months period and the exploration of genome data revealed a limited spectrum of nucleotides changes between the strains within this time lapse. We obtained 5 independent chromosome assemblies using the Pacbio technology which look essentially similar between isolates P1 and P5, thus indicating the robustness of the assembly approaches. Apart from a specific region duplications and chromosome rearrangements (Chr 6), only a few non-synonymous SNPs were identified (17) in addition to a higher number of changes in intergenic regions. Comparisons with other available genomes of *C. lusitaniae* suggested much higher level of nucleotide divergence. When comparing coding regions alone between the *C. lusitaniae* strain ATCC 42720 with P1 to P5, 83225 SNPs were identified. This accounts for a large nucleotide divergence within the same species, a feature also reported in the study of genome comparisons made in other *Candida* spp. (27, 28). Recently, a study reported the analysis of 20 different *C. lusitaniae* isolates from a patient with cystic fibrosis (5). The study reported 404 inter-isolate SNPs (among which 45% were NSS) and 536 indels in total. In comparison, we observed less variation between P1-P5 in the present work. Additional *C. lusitaniae* genomes should be analysed for the further appreciation of genome divergence within this species.

The primary goal of this study was the detailed analysis of genomes between isolates P1 to P5 in order to identify the molecular basis of antifungal resistance in the recovered samples. Our data suggest that each isolate originated from the most susceptible isolate P1. We suggested earlier that the specific drug resistance profiles followed the drug regimen (7). The SNP profiles of each isolate now suggest that each isolate underwent independent micro-evolution trajectories within the host under drug selection. Common SNPs responsible for drug resistance were observed in isolates P3 and P4 (V668G of *MRR1*) and in isolates P2 and P5 (S412* of *ERG4*). This raises the possibility of a closer relationship between these isolate pairs. However, each of the isolates contains additional SNPs and indels which are against the hypothesis of a close lineage, i.e. that each of the P4 and P5 isolates are issued from parents P3 and P2, respectively. Given the number of shared SNPs between the isolates P3/P4 and P2/P5 isolates, it is possible that they originated from common ancestors. We cannot exclude that a more comprehensive isolate collection could have revealed better sequential SNPs relationships between isolates, however, and this is one of the limitations of our study, the isolate sampling strategy was not aimed first to address isolate diversity.

FLC resistance was earlier correlated with the upregulation of the multidrug transporter *MFS7*, a close homolog of the *C. albicans* transporter *MDR1*. Genome comparisons in *C. lusitaniae* isolates P1-P5 identified a SNP in *MRR1* leading to a V668G substitution in P3 and P4. We provided genetic (knock-outs and reversion experiments) to support that the V668G substitution is a GOF mutation which stimulates *MFS7* expression. These data are consistent with a recent study describing several *MRR1* mutations associated with FLC resistance in *C. lusitaniae*, but not including the V668G GOF mutation (5). This mutation may lead to the upregulation of several other genes and this hypothesis was confirmed here by RNAseq analysis of the P3 and P4 isolates. Interestingly, our data matches well with those comparing an azole-resistant with an azole-susceptible *C. lusitaniae* isolate. From the 19 identified regulated genes in this work (5), 12 were corresponding in our data set, and *MDR1* was one among them (File S4).

It was interesting to observe that deletions of *MRR1* and *MFS7* in P3 resulted in different FLC MIC values (MIC=1 µg/mL FLC and 8 µg/mL FLC, respectively). This underscores that, in addition to *MFS7*, *MRR1* controls other FLC resistance mediators. One likely candidate could be the *CDR1* homolog EJF14_30164 which is 2- to 4-fold upregulated in P3 and P4 as compared to P1 (Table 3). *CDR1*, also known as ABC15 (29), was shown previously as slightly upregulated in P3 and P4 as compared to P1 (7). *CDR1* and homologs are known to mediate azole efflux and are important contributor of azole resistance in several fungal pathogens (30). *CDR1* may be controlled by *MRR1* in *C. lusitaniae*. This contrasts with the regulatory properties of *MRR1* in *C. albicans*, which do not include *CDR1* (31). Further work is needed to address this question.

The *MRR1* V668G substitution was associated with another phenotype (resistance to 5-FC) and we propose that the multidrug transporter *MFS7* (a major facilitator efflux transporter) could mediate this phenotype. The data supporting this conclusion are based on genetic approaches as well as *MFS7* overexpression in *C. albicans*. Until now, 5-FC resistance in *Candida* spp. has been looked essentially at the angle of drug import and our work provides evidence that drug efflux may also be involved. One can also argue that the known 5-FC import system (such as *FCY2*) could be regulated in the presence of the *MRR1* V668G mutation, which would result in 5-FC resistance, however our RNAseq data do not support this hypothesis (data not shown). Interestingly, it was shown in *C. lusitaniae* by Noel *et al.* (4) that FLC and 5-FC can exhibit cross-resistance when both drugs are combined *in vitro*, however the mechanism behind this observation remained elusive. In this experimental setting, we propose that *MFS7* could have been upregulated in a transient manner by the addition of both drugs *in vitro*. This possible elevated expression could be sufficient to result in a cross-resistance phenotype. Additional work is warranted to support this hypothesis.

The genome comparisons out of Pacbio assemblies identified the same three separate *FKS1* mutations that were earlier reported in isolates P2, P4 and P5 (7). We also confirmed in our first study that the *FKS1* amino acid substitutions S638P, S638Y and S631Y were causing candin resistance using *S. cerevisiae* as a surrogate (7). Recently, it was reported that S638P substitution could mediate candin resistance in *C. lusitaniae*. This was achieved by allelic replacement in this species (32) and thus confirms the importance of the *FKS1* position 638^Ser^ in establishing candin resistance in *C. lusitaniae*. The Fks1p position 638^Ser^ lies within a region called hot spot region 1 (HS1) that is enriched in amino acid substitutions responsible for candin resistance (32). Position 631^Ser^ is the only position that was yet not been confirmed as causing candin resistance in *C. lusitaniae*, however its proximity to HS1 and our own study with *S. cerevisiae* makes the S631Y substitution likely as a cause of candin resistance.

Amb resistance in fungal pathogens is usually caused by mutations in the ergosterol biosynthesis pathway (33). *C. lusitaniae* develops resistance to Amb during infections and also by inactivation of specific *ERG* genes *in vitro* (34). In this work we provide evidence that Amb resistance is mediated by loss of function of two genes, *ERG3* and *ERG4. ERG3* loss of function mutations have been documented in several fungal pathogens, all resulting in Amb resistance (26, 35–38). To our knowledge, *ERG4* loss of function mutations have not been described so far in fungal pathogens. We show here that *ERG4* loss of function results in absence of ergosterol, thus contributing to Amb resistance as measured earlier (7). A surprising result was the combination of the two loss of function mutations in isolate P5, which was the last recovered isolate from the treated patient. Accumulation of two different loss of function *ERG* mutations in the same isolate is unique to our knowledge in fungal pathogens. Up to now, the combination of *ERG* mutations was only known in *C. albicans* for functional *ERG11* point mutations combined with defects in *ERG5* or *ERG3* leading to both azole and Amb resistance (39, 40). Interestingly, isolate P5 exhibits a slightly higher Amb MIC as compared to P2 (2 vs 1 µg/mL, File S5) and thus it is likely that the accumulation of 2 *ERG* mutations conferred enhanced protection against the activity of Amb in this isolate. Supposing that P2 was the parent of P5, this implies that the *ERG3* mutation occurred sequentially after the emergence of the *ERG4* mutation, however we cannot firmly confirm this hypothesis. The combination of the loss of function mutations, irrespective of their timely occurrence, rather suggests functional compensatory effects. For example, one of the mutations in the *ERG* genes (i.e. *ERG4*) could have resulted in fitness defects and therefore decreased virulence. We addressed this hypothesis by testing the virulence of each isolate in the mini-host *Galleria mellonella*. Our results could not detect significant virulence difference between strains (Fig. S2). The necessity for P5 to carry two independent *ERG3* and *ERG4* loss of function may be therefore solely explained by the resulting benefit in Amb resistance.

Taken together the results of our study highlight that genome comparisons are highly relevant in the resolution of drug resistance mechanisms. Genome sequencing is becoming affordable and several other studies have already taken comparative genomics as a way to detect antifungal resistance mechanisms (41). Nevertheless, the relevance of genome alterations needs to be addressed by parallel experimental approaches that were greatly facilitated by the development of novel genome editing technologies (CRISPR-Cas9).

## Material and Methods

### Strains and media

*C. lusitaniae* and *C. albicans* isolates were grown in complete medium YEPD (1% Bacto peptone, Difco Laboratories, Basel, Switzerland), 0.5% Yeast extract (Difco) and 2% glucose (Fluka, Buchs, Switzerland) at 30 °C under agitation. Plasmids were propagated in *Escherichia coli* DH5α on Luria-Bertani (LB) 2% agar plates at 37°C overnight (42). LB was supplemented with either 100 µg/mL ampicillin (AppliChem) or 34 µg/mL chloramphenicol (Fluka, Buchs, Switzerland) when necessary.

### Antifungal susceptibility testing

Antifungal susceptibility testing was carried out with The Sensititre YeastOne (SYO) colorimetric microdilution method and was performed using commercially available panels (Thermo Fisher Scientific, Switzerland) according to manufacturer’s recommendation. The serial dilutions susceptibility method was performed onto agar plates with either YEPD or YNB minimal medium (0.67% yeast nitrogen base [Dicfo] with 2% glucose. Yeast cells from an overnight culture in YEPD medium were diluted to 1.5 x 10^7^ cells/mL in 1 mL Phosphate Buffer Saline (PBS) (Bischel, Interlaken, Switzerland) and 200 µL of this solution were transferred to a 96-well flat-bottom plate (Sigma-Aldrich). Ten-fold serial dilutions were performed from 1.5 x 10^7^ to 1.5 x 10^2^ cells/mL in PBS. The cell dilutions were next spotted on plates with a 48-pin replicator (V&P Scientific, Inc., San Diego, CA, USA) and plates incubated at 35 °C for 24- to 48 h.

### Pacbio genome sequencing

In order to produce high quality genomic DNA from *C. lusitaniae* isolates, overnight cultures (5 mL) were grown in YNB minimal medium (0.67% yeast nitrogen base [Dicfo] with 2% glucose) at 30°C to obtain 2 x 10^8^ cells for each strain. DNA isolation followed instructions of the Gentra Puregene Yeast/Bact kit (Qiagen) with slight modifications. First, yeast cell lysis was performed with Zymolyase 100T (3 µg/µL end concentration) for 30 min at 37°C. In addition, phenol/chloroform extractions were carried out after RNAse A treatment of precipitated nucleic acids according to recommendations issued by Pacific Bioscience (PacBio SampleNet-shared protocol) for the use of phenol/chloroform/isoamyl alcohol. After final precipitation with NH_4_OAc and several washes with 80% alcohol, the genomic DNA was dissolved carefully in a small volume of elution buffer (10 mM Tris-HCl, pH 8.5). Aliquots of 5 μg of extracted, high-quality, genomic DNA was diluted to 150 μL using elution buffer at 30 μg/μl. Long insert SMRTbell template libraries were prepared (20 kbp insert size) according to PacBio protocols. Each isolate used 2 SMRT cells which were sequenced using P6 polymerase binding and C4 Blue pippin sequencing kits with 240 min acquisition time on PacBio RSII at the Lausanne Genomic Technologies Facility (LGTF) of the University of Lausanne (Unil). De novo genome assemblies were produced using PacBio’s SMRT Portal (v2.3.0) and the hierarchical genome assembly process (HGAP version 3.0), with default settings and a seed read cut-off length of 6000 bp.

### Annotation

For the *C. lusitaniae* annotation process, we took advantage of available RNAseq reads obtained from isolates P1, P3 and P4 isolates (see below). These reads were used to predict and confirm CDS of the sequenced genomes. Processed reads were aligned against the P1 reference genome available as a fasta file from Pacbio assemblies (see below) and the resulting alignments were assembled into potential transcripts using StringTie (v1.3.3) (43). These transcript assemblies were subsequently used as Expressed sequence tags (EST) evidence for assigning the structural annotations to the P1 genome using Work-Queue Maker (WQ-MAKER) (44). Uniprot *C. lusitaniae* protein dataset was downloaded as (https://www.uniprot.org/uniprot/?query=candida+lusitaniae&sort=score). These protein sequences were used as protein evidence during structural annotations. We used MAKER (45) that incorporates SNAP (46) into the gene prediction pipeline to annotate the P1 genome assembly. SNAP gene predictions make use of Hidden Markov Models (HMMs) as their underlying probabilistic model. Repetitive elements, including low-complexity sequences and interspersed repeats in the input genome sequence was identified and masked using RepeatMasker (47) by aligning the non-annotated genome against a library of known repeats such as Repbase (48). Transcript assemblies and protein sequences described above were used as evidence to aid gene predictions. In the initial run, MAKER was launched iteratively and the tasks such as repeat masking and evidence alignments were performed which resulted in GFF3 (general feature format) file containing the masked regions and protein transcript alignments. The GFF3 file generated from the above step was used during subsequent MAKER runs. The data generated in the initial run was used in training the gene predictions using SNAP. All the transcript sequences that were used as evidence during the initial MAKER run were placed into a single transcript fasta file and were used for SNAP HMM training. After training SNAP HMMs from iterative runs that generate imperfect gene models, MAKER was run one again in the final step to accurately predict genes and corresponding ORFs.

Following the detection of ORFs, all proteins of the P1 genome were blasted to the SwissProt database (release-2017_09) using Blast2GO (Version 5.2.5; BioBam Bioinformatics, Valencia, Spain). Top-hits were retrieved from the blast results (E-value cut-off of 10^-3^) and were added to each ORF annotations. ORFs without blast results were re-submitted using Blast2GO to the non-redundant protein NCBI database to enlarge the blast search and positive results were added to the existing annotations.

### Genome comparisons

Annotated genomes were compared using Mauve (version 2015-02-25) with default parameters and aligner software Muscle 3.6. Produced alignments were next imported in Geneious Prime® (2019.1.3, Biomatters, Ltd, Auckland, New Zealand) and each chromosome alignment was exported as a nucleotide alignment file. After establishing genome of isolate P1 as a reference, SNPs as well as insertions/deletions were obtained by the software by selection of coding and non-coding regions.

SNP densities along chromosomes were obtained from VCF files extracted from each chromosome comparisons using Geneious. VCF files corresponding to each chromosome were imported into the online software SNiPlay (49) with a size of the sliding window of 2000 nucleotides. Data were exported in Graph Prism 8.1.2 (GraphPad Software, San Diego, CA, USA) for visualisation.

For P1 to P5 genome comparisons, differences between genomes in homopolymeric regions (poly(dN) stretches of at least 7 nucleotides) were ignored, since they were likely due to artifacts of the Pacbio sequencing technology as reported earlier (50). Only differences in these stretches occurring in at least 2 separate genomes were considered in genome comparisons. In addition, only telomeric chromosome regions that were common to all P1 to P5 isolates were included in genome comparisons.

### Genome-wide transcriptional analysis

#### (i) RNA extraction and processing

RNA was isolated from 5 mL log phase cultures of isolates P1, P3 and P4 grown in YEPD medium at 30°C under agitation. Total RNA was extracted from with the RNeasy Protect Mini kit (Qiagen) by a process involving mechanical disruption of the cells with glass beads as previously described (51). Total RNA extracts were treated with DNase using a DNA-free kit (Ambion-Life Technologies, Zug, Switzerland). RNA quality and integrity were verified with a fragment analyzer automated CE system (Advanced Analytical). RNA extractions were performed in biological triplicates. RNA libraries for RNAseq were prepared with a TruSeq Stranded Total RNA Library Prep Kit (Illumina). The resulting libraries were sequenced on an Illumina HiSeq 2500 system at the LGTF.

#### (ii) Quality control and processing of raw RNAseq reads

Twenty-seven single end RNAseq datasets from three different *C. lusitaniae* strains (P1, P3 and P4) after RNA sequencing were obtained. Quality of the raw reads were assessed with FastQC (v0.11.7) (52). Prior to mapping and assembly, adapter trimming, quality filtering, artefact removal and contaminant filtering were carried out with BBDuk (v38.51) from BBTools package (53). Low complexity filtering and removal of rRNA sequences were carried out using Prinseq (v0.20.3) (54) and SortMeRNA (v2.1) (55), respectively. Processed reads were aligned to the P1 annotated genome using HSAT2 (v2.1) (56).

#### (iii) Differential Gene expression analysis

Data normalization and differential expression analysis were performed in R (v3.5.2) using Bioconductor packages. The read count data were normalized with the TMM (trimmed mean of M-values) method available in the R Bioconductor package edgeR (57) and subsequently transformed to log2 counts per million by Voom, a method implemented in the R Bioconductor package Limma (58). A linear model with one factor per condition was applied to the transformed data using Limma (59).

### *MRR1*/*MFS7* gene deletions

#### Plasmid constructions

In order to delete *MRR1* and *MFS7* in *C. lusitaniae*, flanking regions of the two genes were cloned by PCR into pSFS2A (60). *MRR1* flanking regions were amplified from isolate P1 with primers pairs ClMRR1-Apa/ClMRR1-Xho and ClMRR-SacI/ClMRR-SacII (Table S1), respectively. *MFS7* flanking regions were amplified from isolate P1 with primers pairs MFS7-Kpn/MSF7-Xho and MFS7-SacI/MFS7-SacII (Table S1), respectively. PCR products were cloned sequentially into corresponding sites to result in plasmid pDS1860 (*MFS7* inactivation) and pDS1864 (*MRR1* inactivation). pDS1860 and pDS1864 were next modified by removing the *SAT1* flipper cassette system by *Bam*HI/*Not*I digestions and replacement with the *NAT1* dominant marker amplified from pJK795 (61) using primers NAT1-BglII and NAT1-Not, thus resulting in pDS2039 and pSD2038, respectively.

#### Cas9-CRISPR for knock-out of *MRR1* and *MFS7*

The RNA-protein complexes (RNPs) approach reported in Grahl *et al.* (62) was used that employs reconstituted purified Cas9 protein in complex with scaffold and gene-specific guide RNAs. gRNA specific for *MRR1* and *MFS7* (Table S1) were selected and obtained from IDT (Integrated DNA Technologies, Inc.) as CRISPR guide RNA (crRNA), which contains 20-bp homologous to the target gene fused to the scaffold sequence. Gene-specific RNA guides were designed in silico using Geneious Prime. RNPs were created using the Alt-R CRISPR-Cas9 system from IDT. Briefly, crRNAs (crMRR1 and crMFS7, Table S2) and tracrRNA (a universal transactivating CRISPR RNA) were first dissolved in RNase-free distilled water (dH_2_O) at 100 µM and stored at −80°C. The complete guide RNA was generated by mixing equimolar concentrations (4 µM final) of the gene-specific crRNA and tracrRNA to obtain a final volume of 3.6 µl per transformation. The mix was incubated at 95°C for 5 min and cool down to room temperature. The Cas9 nuclease 3NLS (60 µM stock from IDT) was diluted to 4 µM in dH_2_O at a volume of 3 µl per transformation. RNPs were assembled by mixing guide RNAs (3.6 µl of gene-specific crRNA/tracrRNA) with 3 µl of diluted Cas9 protein, followed by incubation at room temperature for 5 min. Transformation of *C. lusitaniae* cells was carried out by electroporation and used 6.6 µl of gene-specific RNPs, 40 µl of *C. lusitaniae* cells and 1-2 µg of repair constructs (up to 3.4 µl volume). Repair constructs containing the *MRR1* and *MFS7* inactivation cassettes were obtained by PCR amplification with primer pairs ClMRR1-Apa/ClMRR-SacI and MFS7-Kpn/MFS7-SacI from pDS2038 and pSD2039, respectively. Transformants were selected at 30 °C on YEPD agar containing 200 µg/mL nourseothricin. Transformants were verified by PCR using primer pairs NAT1_134_R/ClMRR1-verif3 for *MRR1* deletion and NAT1_134_R/5-MFS7-A for *MFS7* deletion. DNA from transformants was prepared by small-scale rapid DNA extraction as described (63).

#### *MRR1* reversion

In order to re-introduce *MRR1* alleles in the background of *MRR1* deletion mutants, an alternative mutant construction using a recyclable *NAT1* marker was employed. The maltose-inducible *MAL2*-*FLP1* system of pSFS2A was first excised from pSFS2A as a 0.9 kb *Apa*I-*Eco*RV fragment and cloned into pJK863 to substitute the *SAP2* promoter, thus resulting in pDS2046. In this approach, the *NAT1* marker could be recycled by *MAL2*-dependent *FLP1* expression (MAL2-FLP-NAT1). This plasmid was used as a PCR template with the primer pair MRR1-5_pDS2046/MRR1-3_pDS2046. Both primers contained 70-bp homology to the *MRR1* 5’- and 3’-flanking regions and a 21-bp extension matching to the MAL2-FLP-NAT1 extremities. CRISPR-Cas9-mediated recombinations at *MRR1* flanking regions with this PCR-amplified repair fragment was used with crRNAs crMRR1_del5 and crMRR1_del3 that were prepared as explained above to reconstitute functional RNPs, with the exception that both RNPs were concentrated by 2-fold. Transformation of *C. lusitaniae* was carried out by electroporation as described below and transformants selected onto YEPD plates with nourseothricin (200 µg/mL).

After *MRR1* deletion verification by PCR as described above, strains were grown overnight on YEP liquid medium with 2% maltose at 30 °C in order to induce *FLP1*-mediated recombination at FRT sequences and resulting loss of *NAT1*. Recycling of *NAT1* was observed in YEPD agar medium containing each about 10^2^ *C. lusitaniae* cells at a nourseothricin concentration of 1 µg/mL to distinguish between parent cells and those without *NAT1*.

Nourseothricin-sensitive *C. lusitaniae* cells deleted for *MRR1* were used for *MRR1* reversion. *MRR1* alleles were first cloned into pSD2038 with fragments obtained by PCR using primers ClMRR1-Apa and ClMRR1-Xhorev and DNA templates from isolate P1 and P3, which resulted in plasmids pDS2040 and pDS2041, respectively. Repair fragments were obtained from both plasmids with primers ClMRR1-Apa and MRR1-3 rev new. CRISPR-Cas9-mediated recombinations at *MRR1* flanking regions with these PCR-amplified repair fragments were used with crRNAs crRNA_MRR1_rev5 et crRNA_MRR1_rev3 that were prepared as above-explained to reconstitute functional RNPs. Transformations of *C. lusitaniae* were carried out by electroporation as described below and transformants selected onto YEPD plates with nourseothricin (200 µg/mL). Re-integration of *MRR1* alleles was verified by PCR on recovered genomic DNA with primer pair ClMRR1_F/ ClMRR1_3377_R followed by sequencing with primer ClMRR1_2900_F to confirm allele identity.

### *MFS7*-GFP tagging

GFP tagging of *MFS7* was realized by a Gateway cloning approach developed by Chauvel *et al.* (64). *MFS7* was first amplified with primers Forward_gateway_MFS7 Reverse_gateway_MFS7 (Table S1) and cloned into pDONR207 (64) by recombination reaction with Invitrogen Gateway BP Clonase™. Next, *MFS7* was transferred into pCA-DEST1102 (a plasmid aimed to express C-terminal GFP fusion proteins under the control of a tetracycline-regulated promoter) by LR Clonase™ to result into pDS2034. Next, this plasmid was linearized by i-*Sce*I and transformed for Ura^+^ selection in strain CPY41, a derivative of SC5314 containing pNIMX (64) and lacking *URA3* alleles, to yield DSY5194.

### *MFS7* overexpression

*MFS7* overexpression used a system that overexpresses proteins in *C. albicans* through a *CDR1* promoter under the control of a *TAC1* gain of function allele (25). The *MFS7* ORF was amplified from isolate P1 with primers MFS7_XbaI et MFS7_Nhe and cloned into the single *Spe*I site of pDS1873, a derivative of pDS1874 lacking *CDR1-GFP* (25) to yield pDS2022. This plasmid was digested by *Sac*I et *Pvu*I to allow recombination at the *CDR1* locus and transformed into DSY4684 lacking major multidrug transporters (25). As control, DSY4684 was transformed with CIp10 after *Stu*I digestion (65). Transformants were obtained by Ura+ selection.

### *ERG3* and *ERG4* reversions

In order to restore wild type copies of *ERG3* and *ERG4* in isolates with defects in these genes, a CRISPR approach was used. First, repair fragments were constructed by fusion PCR. The *ERG4* repair fragment was generated by fusion of three PCR fragments with overlapping sequences. The first PCR fragment amplified the wild type *ERG4* from isolate P1 with primers ERG4-P1 and ERG4-NAT1-R (20-bp overlap with pJK795 (61)). The second PCR fragment amplified the selective marker *NAT1* from pJK795 with primers ERG4-NAT1-F (20-bp overlap with *ERG4*) and ERG4-NAT1t-R (20-bp overlap with pJK795). The third PCR fragment amplified a 3’-end portion of *ERG4* with primers ERG3-P2 x ERG4-NAT1t-F (20-bp overlap with pJK795). The final PCR was performed with the 3 purified fragments and nested primers ERG4-P3 and ERG4-P4 in the presence of 1.2 M betaine. The *ERG3* repair fragment was constructed in a similar fashion but using the hygromycin resistance selection marker from pYM70 (66). The first PCR fragment amplified the wild type *ERG3* from isolate P1 with primers ERG3-P1 and ERG3-HYG-R (20-bp overlap with pYM70). The second PCR fragment amplified the selective marker *CaHygB* from pYM70 with primers ERG3-HYG-F (20-bp overlap with *ERG3*) and ERG3-HYGt-R (20-bp overlap with pYM70). The third PCR fragment amplified a 3’-end portion of *ERG3* with primers ERG3-P2 x ERG3-HYGt-F (20-bp overlap with pJK795). The final PCR was performed with the 3 purified fragments and nested primers ERG3-P3 and ERG3-P4 in the presence of 1.2 M betaine.

RNPs were reconstituted as described above. gRNA specific for *ERG3* and *ERG4* (Table S1) were selected for targeting the mutated positions in isolates P2 and P5 and were obtained from IDT (Integrated DNA Technologies, Inc.) as CRISPR guide RNA (crRNA) fused to the scaffold sequence. RNPs were assembled by mixing guide RNAs (3.6 µl of gene-specific crRNA/tracrRNA) with 3 µl of diluted Cas9 protein, followed by incubation at room temperature for 5 min. Transformation of *C. lusitaniae* cells was carried out by electroporation as described below and used 6.6 µl of gene-specific RNPs, 40 µl of *C. lusitaniae* cells and 1-2 µg of *ERG3* or *ERG4* repair constructs (up to 3.4 µl volume). Isolates P2 and P5 were used with the *ERG4* repair fragment and transformants selection was operated onto nourseothricin selective medium (200 µg/mL nourseothricin). The *ERG3* repair fragment was used with an *ERG4* revertant from P5 and transformants selection was operated onto hygromycin selective medium (250 µg/mL hygromycin).

Verification of *ERG3* and *ERG4* reversions was carried out by PCR with primers ClERG4_1427_F and NAT1_134_R (*ERG4*) and primers ClERG3_1,104_F and ERG3-NAT1-R (*ERG3*) followed by sequencing.

### *C. lusitaniae* transformation

Transformations in *C. lusitaniae* were carried out by electroporation. Overnight-grown cells were freshly diluted 20-fold in 20 mL YEPD medium and grown to logarithmic phase to a density of 1-2 10^7^ cells/mL at 30°C under constant agitation. After centrifugation for 5 min at 2500 rpm at 4°C and removal of supernatant, cells were resuspended in 10 mL transformation buffer (100 mM LiAc, 100 mM DTT, 10 mM Tris-HCl 7.5, 1 mM EDTA) and incubated for 1 h at room temperature under mild shaking. Cells were next washed twice with ice-cold water and with 1 M sorbitol. Cell pellets were finally resuspended in 200 µl. Transformations were carried out with aliquots of 40 µl slurry in 0.2 electroporation cuvettes (Biorad) with the following settings: 1.8 kV, 200 Ω, 25 µF. After electroporation pulse, 1 mL YEPD was immediately added and cells were incubated overnight with gentle shaking at 30°C. Once the cells were allowed to recover overnight at room temperature, aliquots were plated onto the corresponding selective media.

### Quantitative reverse transcription PCR (qRT-PCR)

Total RNA was extracted as described above from log phase cultures grown in YEPD at 30°C under constant agitation. Gene expression levels were determined by real-time qRT-PCR in a StepOne Real-time PCR System (Applied Biosystems) using the Mesa Blue qPCR Mastermix Plus for Sybr assay kit (Eurogentec). Each reaction was performed in triplicate on three separate occasions. Expression levels of *MFS7* was normalized by *ACT1* expression as described previously (7).

### Sterol analysis

Overnight cultures of *C. lusitaniae* strains were used to inoculate 10 mL YEPD at a starting concentration of 1 x 10^4^ cells/mL. Cultures were grown for 18 h at 37°C and 200 rpm. Cells were then harvested and pellets washed twice with ddH_2_O. Sterols were extracted and derivatised as previously described (67). Briefly, lipids were saponified using alcoholic KOH and non-saponifiable lipids extracted with hexane. Samples were dried in a vacuum centrifuge and were derivatised by the addition of 0.1mL BSTFA (N,O-Bis(trimethylsilyl)trifluoroacetamide) TMCS (Trimethylchlorosilane) (99:1, Sigma) and 0.3 mL anhydrous pyridine (Sigma) and heated at 80°C for 2 hours. TMS-derivatised sterols were analysed and identified using GC/MS (Thermo 1300 GC coupled to a Thermo ISQ mass spectrometer, Thermo Scientific) and Xcalibur software (Thermo Scientific). The retention times and fragmentation spectra for known standards were used to identify sterols.

### Virulence assays with *Galleria mellonella*

*Galleria mellonella* larvae were purchased from TruLarvTM (Biosystems Technology, Exeter, Devon, UK). Larvae weighing between 0.2-0.3g were selected for our experiments and stored at 16°C upon arrival. A total of 10 larvae were infected with each isolate by microinjection (Omnican®100, BBraun). *C. lusitaniae* cells were grown overnight in YEPD medium at 30°C under agitation and pelleted by centrifugation (5 min at 4600 rpm). Cells were washed twice with phosphate buffered saline (PBS) (137 mM NaCl, 10 mM Phosphate, 2.7 mM KCl, and pH 7.4) and the amount of cells was estimated with a photometer (Novaspec® II, Pharmacia). Forty µl of a cell suspension containing 1,5 x 10^7^ cells/mL in PBS was then injected into the last left proleg. Control larvae were injected with the same volume of PBS. The injected larvae were incubated at 30°C in the dark for 9 days post infection and survival was scored each day. Larvae were scored as dead when melanized and upon lack of response after gentle manipulation with a clamp.

### Microscopy

Epifluorescence and phase-contrast imaging were performed with a Zeiss Axioplan 2 microscope equipped for epifluorescence microscopy with a 100-W mercury high-pressure bulb and Zeiss filter set 9 (for GFP imaging). Images obtained with a SPOT RT3 cooled 2 Mp CCD camera (Diagnotic Instruments Inc., MI, USA) were recorded and captured with VisiView (Visitron Systems GmbH, Germany).

### Data availability

Strains described here are available upon request. PacBio assemblies are available under Bioproject PRJNA504391. RNAseq data are available under study SRP172837. The final assembled genomes for isolates P1 to P5 and their annotations are available under accession numbers CP038484-CP038491, CP039550-CP039557, CP039652-CP039659, CP039618-CP039625 and CP039660-CP039667, respectively.

## Acknowledgements

The authors are thankful to Danielle Brandalise and Francoise Ischer for excellent technical assistance. The authors are also thankful to Emmanuel Beaudoing for discussions on genome sequencing approaches. The *C. albicans* Gateway plasmids were kindly provided by C. d’Enfert (Pasteur Institute, Paris). This study was partially financed by a Swiss Research National Foundation grant 31003A_172958 to DS.

## Supplementary material

**Table S1:** Primers and guides used in this study

**Table S2:** Strains used in this study

**Table S3:** Chromosome lengths of *C. lusitianiae* chomosomes

**Figure S1:** SNP density plots from comparisons between *C. lusitaniae* isolate P1 and ATCC 42720. Red bars correspond to the lengths of each indicated chromosome.

**Figure S2:** *Galleria mellonella* infection with isolates P1 to P5. Data were plotted with Graph Prism 8.1.2 (GraphPad Software, San Diego, CA, USA) and no statistical differences were noticed between the survival of the tested isolates.

**File S1:** SNP counts between ATCC42720 and isolate P1

**File S2:** Transposons and LTRs of *C. lusitaniae* and *GRP2* homologs in *C. lusitaniae*

**File S3:** Nucleotide differences between isolates P1 to P5

**File S4:** RNAseq data comparisons between isolates P3 and P4 versus isolate P1

**File S5:** Antifungal susceptibility data

